# Costimulatory domains direct distinct fates of CAR-driven T cell dysfunction

**DOI:** 10.1101/2023.01.26.525725

**Authors:** Mehmet Emrah Selli, Jack H. Landmann, Marina Terekhova, John Lattin, Amanda Heard, Yu-Sung Hsu, Tien-Ching Chang, Jufang Chang, John Warrington, Helen Ha, Natalie Kingston, Graham Hogg, Michael Slade, Melissa M. Berrien-Elliot, Mark Foster, Samantha Kersting-Schadek, Agata Gruszczynska, David DeNardo, Todd A. Fehniger, Maxim Artyomov, Nathan Singh

## Abstract

Chimeric antigen receptor (CAR) engineered T cells often fail to enact effector functions after infusion into patients. Understanding the biological pathways that lead CAR T cells to failure is of critical importance in the design of more effective therapies. We developed and validated an *in vitro* model that drives T cell dysfunction through chronic CAR activation and interrogated how CAR costimulatory domains contribute to T cell failure. We found that dysfunctional CD28-based CARs targeting CD19 bear hallmarks of classical T cell exhaustion while dysfunctional 41BB-based CARs are phenotypically, transcriptionally and epigenetically distinct. We confirmed activation of this unique transcriptional program in CAR T cells that failed to control clinical disease. Further, we demonstrate that 41BB-dependent activation of the transcription factor FOXO3 is a significant contributor to this dysfunction and disruption of *FOXO3* improves CAR T cell function. These findings identify that chronic activation of 41BB leads to novel state of T cell dysfunction that can be alleviated by genetic modification of FOXO3.

**Summary:** Chronic stimulation of CARs containing the 41BB costimulatory domain leads to a novel state of T cell dysfunction that is distinct from T cell exhaustion.

## Introduction

A growing body of literature suggests that failure of chimeric antigen receptor (CAR)-engineered T cells against both hematologic and solid cancers most often results from defects in normal T cell function (*1*). These impairments manifest as poor CAR T cell expansion, limited persistence and ineffective anti-tumor cytotoxicity. Correlative studies have identified an association between high tumor burden and limited anti-tumor activity, linking prolonged or high-intensity receptor activation with CAR T cell failure (*2-4*). These limitations resemble the functional defects seen in exhaustion, a cellular state that develops from persistent stimulation of the native T cell receptor (TCR) in the setting of chronic viral infection, cancer and autoimmunity (*5*). Several recent pre-clinical studies have confirmed that persistent CAR activation results in the acquisition of a dysfunctional T cell state that shares many functional attributes of exhaustion (*6-9*). Collectively, these findings clearly indicate that chronic antigen receptor activation is detrimental for both TCR and CAR-driven T cell function, and the prevailing presumption is that CAR T cells fail in patients because they become exhausted.

CAR design integrates activating signals from the TCR and a required costimulatory receptor. Of the six FDA-approved CAR T cell products, two contain the signaling domain from CD28, the paradigmatic costimulator with a central role in the endogenous T cell response to infection. The four other products contain the signaling domain from 41BB, a member of the tumor necrosis factor receptor (TNFR) superfamily with a narrower role in T cell activation primarily described in the context of memory T cell development. A plethora of pre-clinical and clinical data confirm that these CAR-integrated costimulatory domains have a significant impact on function, most notably directing distinct patterns of CAR T cell expansion, persistence and therapy-related toxicity (*10*). Consideration of CAR product costimulatory domains is central to clinical decision making, however how costimulatory domains contribute to the T cell dysfunction that is often responsible for clinical failure has not been investigated.

To interrogate this central question, we developed an *in vitro* system to chronically activate CD19-directed CAR T cells with CD19+ acute lymphoblastic leukemia (ALL) cells (*11*). We observed similar functional defects for both CD28 and 41BB-based CAR T cells, but divergent transcriptional, epigenetic and phenotypic attributes. While CD28-based CAR T cells bore the hallmarks of T cell exhaustion, 41BB-based cells activated a distinct molecular program not previously associated with T cell dysfunction. Evaluation of CAR T cells collected several months after treatment from a patient with progressive lymphoma confirmed expression of this gene signature in an actively failing clinical product. Through integrated epigenetic and transcriptional analysis, we found that the transcription factor FOXO3 was a driver of 41BB but not CD28-based CAR T cell dysfunction, and that disruption of *FOXO3* alleviated dysfunction *in vitro* and *in vivo*. Together, these data identify a unique CAR-driven dysfunction program activated by 41BB and regulated by FOXO3.

## Results

### Persistent exposure to CD19+ tumors drives dysfunction of CD28 and 41BB-based CAR T cells

In order to understand how prolonged costimulatory domain activation impacts CAR T cell function, we developed an *in vitro* model in which CD19-directed human CAR T cells containing either CD28 (19/28) or 41BB (19/BB) domains were combined with the CD19+ ALL cell line Nalm6. These co-cultures were established at an effector-to-target (E:T) ratio of 1:8 and monitored for dynamic changes in quantity of CAR T cells and Nalm6. As CAR T cells eliminated Nalm6 and proliferated in response to activation we maintained consistent CAR stimulation by replenishing cultures with additional Nalm6 to re-establish an E:T ratio of 1:8 every ∼48 hours (**Figure 1a**) (*11*). After initial engineering with the CAR lentiviral constructs, T cell products were purified using magnetic beads targeting a truncated CD34 selection marker (encoded on the CAR-containing plasmid but separated from the CAR transgene by a P2A sequence) to ensure all T cells in co-cultures were CAR+. We observed comparable CAR expression after purification as well as at the end of chronic stimulation co-cultures (**Extended Data Fig. 1a**). Longitudinal measurement of T cell proliferation, a core attribute of effective T cell function, revealed robust expansion of both cell types with greater expansion of 19/28 than 19/BB (**Figure 1b**), as has been observed with clinical products targeting CD19 (*10*). After ∼13 days of chronic stimulation, however, both 19/28 and 19/BB lost proliferative capacity and contracted, reflecting loss of function. Since cumulative Nalm6 survival could not be longitudinally measured in this re-feeding model, we used E:T ratio over time as measure of cytotoxic function. 19/28 and 19/BB demonstrated robust cytotoxicity against persistently high tumor burdens until ∼day 13, at which time both cell types lost the ability to kill (**Figure 1c**). 19/28 and 19/BB isolated after 7 days of chronic stimulation re-stimulated with fresh Nalm6, which led to T cell activation and potent secretion of effector cytokines IFNγ and TNFα (**Figures 1d-e**). In contrast, cells collected after 15 days of chronic stimulation secreted almost no cytokine in response to re-stimulation. We repeated these assays using T cells derived from four independent donors and found no difference in the timing of dysfunction onset (**Figure 1f**), defined as either expansion of Nalm6, reflecting a failure to control disease, or decline in T cell count, reflecting failure of T cell expansion.

**Figure 1.**
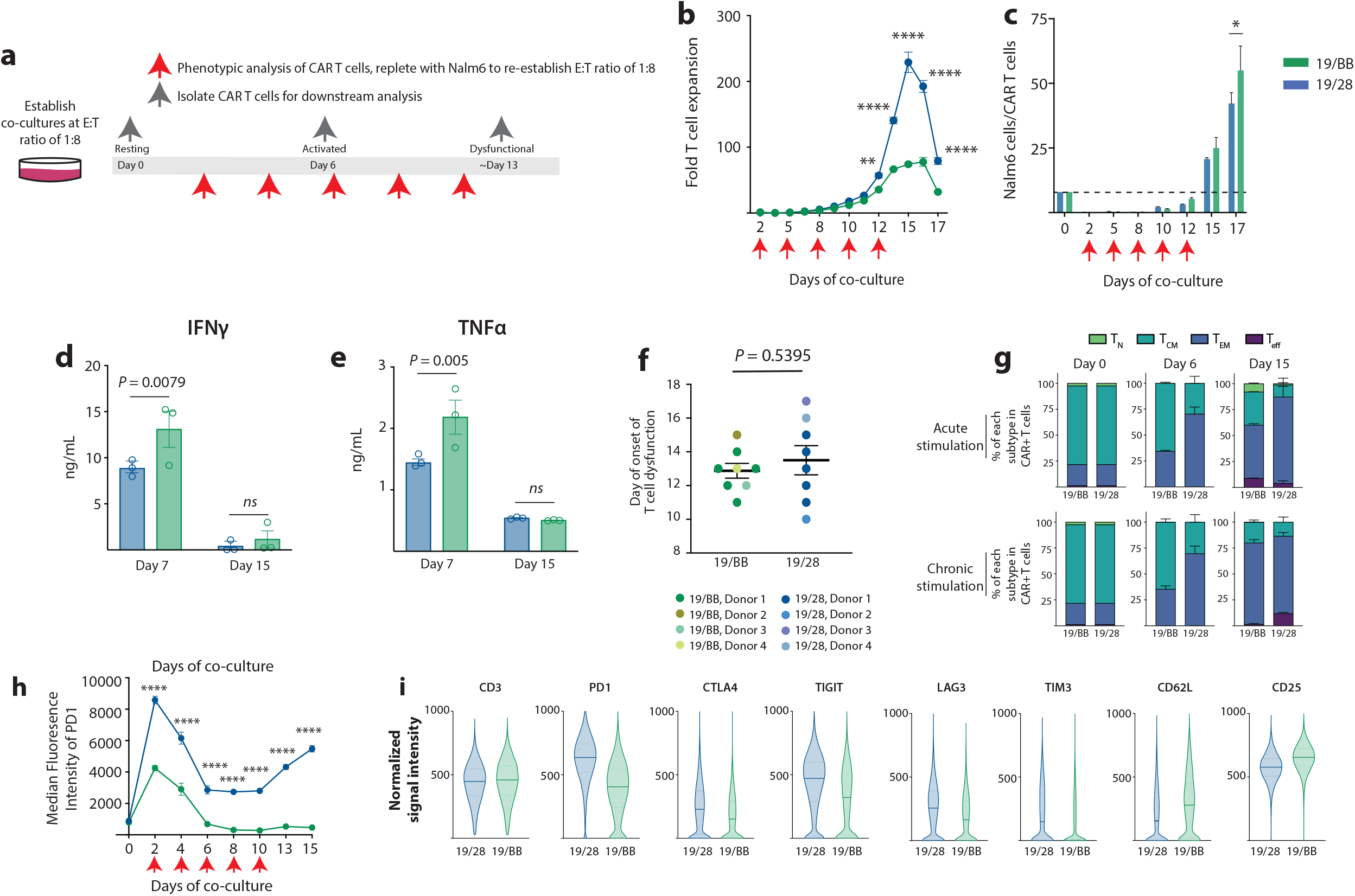
Chronic CAR stimulation results in T cell dysfunction. **a**, Schematic of *in vitro* chronic stimulation assay. **b**, Expansion of 19/28 and 19/BB CAR T cells over the course of chronic stimulation. **c**, target Nalm6 cells per CAR T cells over the course of chronic stimulation. **d-e**, Production of **d**, INFγ and **e**, TNFα by 19/28 and 19/BB cells isolated at days 7 and 15 of chronic stimulation cultures upon re-stimulation. **f**, Kinetics of dysfunction onset as reflected by first day of failure as measured by T cell contraction or loss of tumor control. Data from n=4 independent donors. **g**, Change in memory phenotype of CD8+ CAR T cell products after either acute (single combination with Nalm6 cells) or chronic stimulation. **h**, Surface expression of PD1 during chronic stimulation. **i**, Violin plots of T cell markers on day 15 cells evaluated by CyTOF. **b-e, g-h** are representative data from n=4 independent donors, **i**, representative of n=2 donors. * P<0.05, **P<0.01, ***P<0.001, ****P<0.0001 by two-tailed ANOVA with Bonferroni correction for multiple comparisons.

To shed light on any distinctions in cell state that may develop as a result of persistent costimulation we performed phenotypic analyses of CAR T cells over time. Flow cytometry-based assessment of T cell memory markers revealed few distinctions in CD8 cells as a result of costimulation and confirmed previous observations that 41BB promotes earlier-lineage memory phenotypes (*12*) (**Figure 1g**). This was most pronounced in the setting of acute stimulation, in which co-cultures were established and serially monitored but never replenished with Nalm6 and thus only stimulated for a short time (∼3-5 days). A similar trajectory was observed for CD4 T cells (**Extended Data Fig. 1b**). We next used Jurkat cells, an immortalized T cell leukemia line, engineered with reporters that express distinct fluorescent proteins when NFAT, NFκB and AP1 activate, to assess changes in transcription factor regulation. Chronic stimulation of 19/28 Jurkats resulted in increased NFAT and AP1 activity while 19/BB Jurkats had higher NFκB activity (**Extended Data Fig. 1c**). Persistent activation of NFAT and AP1 has previously been implicated in the development of exhaustion in models of chronic viral infection (*13*) and cancer (*8, 14*). NFκB, a known target of 41BB (*15*) has, to our knowledge, not been shown to be a driver of T cell dysfunction. To determine if these two dysfunctional T cell populations bore the phenotypic hallmarks of exhaustion we evaluated expression of PD1 on CAR T cells over time. While both 19/28 and 19/BB cells demonstrated an initial increase in PD1 expression, reflecting T cell activation, resurgent PD1 expression was only seen in 19/28 cells (**Figure 1h**). We further probed the evolution of T cell phenotypes using an antibody panel targeting 39 intra- and extracellular proteins (**Supplementary Table 1**). Cytometry by time of flight (CyTOF) analysis revealed that pre-stimulation (day 0) and peak-stimulation (day 6) 19/28 and 19/BB cells had very similar expression of these markers, however dysfunctional (day 15) samples occupied slightly distinct spaces by nearest neighbor (tSNE) analysis (**Extended Data Figs. 1d-e**). Dysfunctional 19/28 cells expressed significantly higher levels of known exhaustion markers PD1 and TIGIT while dysfunctional 19/BB cells expressed higher levels of the memory marker CD62L (*SELL*, **Extended Data Figs. 1f-h**). Consistent with these findings, we found that dysfunctional 19/28 cells expressed higher levels of exhaustion markers CTLA4, LAG3 and TIM3 as well, whereas 19/BB expressed higher levels of CD25 (**Figure 1i**). Together, we observed consistently higher expression of known exhaustion markers in 19/28 as compared to 19/BB.

### Dysfunctional CD28 and 41BB-based CAR T cells are transcriptionally and epigenetically distinct

To interrogate the gene expression programs associated with these distinct phenotypic states we performed RNA sequencing of purified CAR T cells at days 0 (n=1 donor), 6 and 15 (n=4 donors for both time points) of chronic stimulation. Principal component analysis (PCA) demonstrated minimal differences between 19/28 and 19/BB cells at days 0 and 6, but a divergence at day 15 (**Figure 2a**). Intriguingly, while day 6 cells segregated from day 0 cells on both PC1 and PC2, day 15 cells were similar to day 0 cells on PC1, suggesting a regression to a resting-like state. Pooled analysis of sequencing from all four donors demonstrated no differentially expressed genes (DEGs) at day 0 (filtered for log_2_-fold change >1.5 and *FDR* <0.05, **Extended Data Fig. 2a**) and only 60 DEGs at day 6 (**Extended Data Fig. 2b**). After the onset of dysfunction (day 15), however, we identified 365 DEGs that segregated into six clusters (**Figure 2b**). Clusters with higher expression in day 15 19/28 cells were enriched for known exhaustion-associated genes (*NR4A2, LAG3, PDCD1, MAF, BATF, CTLA4, NR4A1*). Clusters with higher expression in day 15 19/BB cells were instead enriched for memory markers (*TCF7, IL7R, LEF1*), class II HLA genes (*HLA-DMB, DQB1, DRB1*) (*16*) and genes not previously associated with T cell dysfunction (*TEC, AXL*). Visualization by volcano plot underscores the significantly greater expression of many exhaustion-associated genes in 19/28 cells as compared to 19/BB cells (**Figure 2c**). Tracking expression of characteristic exhaustion genes *PDCD1, CTLA4, LAG3* and *HAVCR2* (encoding TIM3) over time demonstrated a significant rise in expression from day 7 to day 15 in 19/28 cells, as is observed in classical exhaustion, whereas expression of these markers in 19/BB was only mildly increased as cells progressed to dysfunction (**Figure 2d**). Pathway enrichment analysis of DEGs at day 15 demonstrated some overlap but also enrichment of some unique pathways, including TCR signaling, JAK-STAT signaling, PDL1 and PD1 expression pathways in 19/28 cells and graft-versus-host disease and antigen processing and presentation in 19/BB cells (**Extended Data Fig. 2c**).

**Figure 2.**
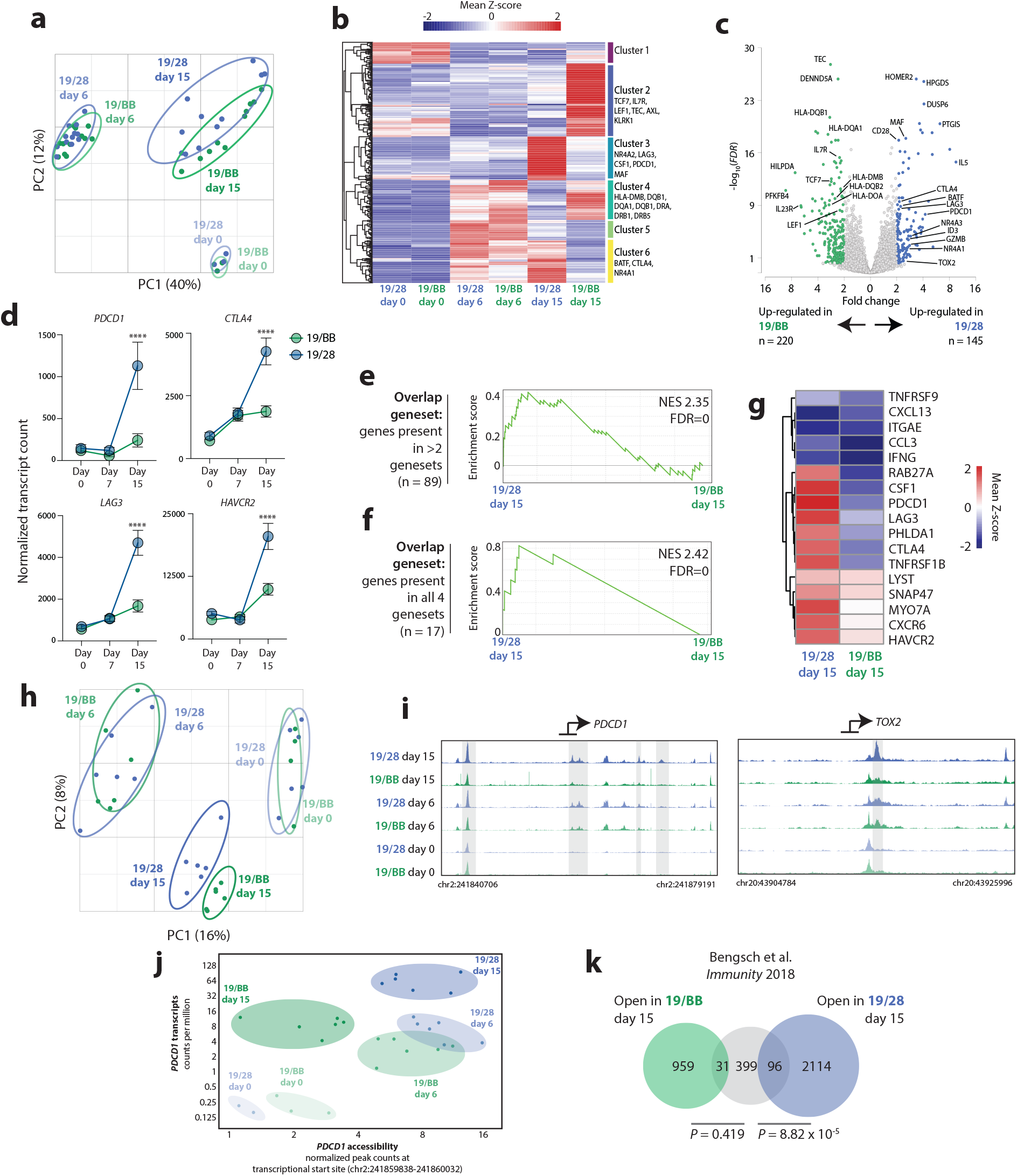
Chronic activation of costimulatory domains directs distinct genomic activity. **a**, Principal Component Analysis (PCA) of bulk RNA sequencing of 19/28 and 19/BB cells at days 0, 6 and 15. **b**, Heatmap of differentially expressed genes between day 15 19/28 and 19/BB cells. **c**, Volcano plot of day 15 differentially expressed genes. n=1 donor for day 0 samples and n=4 donors for days 6 and 15. **d**, Normalized transcript counts of key exhaustion markers over time. Significance determined using two-way ANOVA. **e-f**, Geneset enrichment analysis (GSEA) of genes present in **d**, >2/4 and **f**, all 4 human TIL exhaustion genesets. **g**, Heatmap of day 15 sample expression of genes present in all 4 human TIL exhaustion genesets. **h**, PCA of ATAC sequencing of 19/28 and 19/BB cells at days 0, 6 and 15. n=2 donors for all time points. **i**, Gene tracks at transcriptional start sites of *PDCD1* and *TOX2* reflecting chromatin accessibility. **j**, Correlated transcript count and transcriptional start site accessibility for *PDCD1* over time. **k**, Overlap of genes with increased accessibility in exhausted T cells (as defined in Ref. 22) and genes with increased accessibility in dysfunctional 19/28 or 19/BB cells.

Given the distinction in expression of exhaustion-associated genes, we sought to quantify the similarity of both dysfunctional CAR T cell states to exhausted T cells. Much of the work that has defined the mechanisms leading to exhaustion has been done using murine models of chronic lymphocytic choriomeningitis virus (LCMV). To interrogate genes associated with human T cell exhaustion we turned to four recent studies that profiled gene expression in exhausted tumor-infiltrating lymphocytes (TILs) (*17-20*). We found significantly greater overlap between dysfunctional 19/28 cells and all human exhaustion genesets than we did for dysfunctional 19/BB cells (**Extended Data Figs. 2d-g**). We cross-referenced these TIL genesets to identify genes shared by at least two (n=89) and all four (“master exhaustion” geneset, n=17, **Supplementary Table 2**). Geneset enrichment analysis (GSEA) demonstrated significant enrichment of both shared genesets in dysfunctional 19/28 as compared to dysfunctional 19/BB **(Figures 2e-f**) and higher overall expression of nearly all 17 master exhaustion genes in dysfunctional 19/28 (**Figure 2g**), reflecting similarity in transcriptional identity between classically exhausted T cells and dysfunctional 19/28 but not dysfunctional 19/BB.

In addition to transcriptional identity, several studies have demonstrated recurrent epigenetic alterations that define T cell exhaustion (*21, 22*). To probe how chromatin accessibility evolved over the course of chronic CAR stimulation, we performed longitudinal assay for transposase accessible chromatin with sequencing (ATACseq). Highly consistent with our transcriptional data, we found a divergence in chromatin accessibility only after the onset of dysfunction at day 15, and again observed that day 15 cells drifted back towards day 0 cells on PC1 (**Figure 2h**). Interrogation of two specific exhaustion-associated genes *PDCD1* and *TOX2* revealed significantly greater accessibility at transcriptional start sites in dysfunctional 19/28 as compared to all other groups (**Figure 2i**). We focused on *PDCD1*, an archetypal marker of exhaustion and the target of many therapies that aim to reinvigorate exhausted T cell function, and simultaneously traced chromatin accessibility and transcript counts to profile gene regulatory dynamics at this site. 19/28 cells increased *PDCD1* expression over time and preserved chromatin accessibility as they progressed from activated to dysfunctional, as do exhausted T cells (*21, 22*). In contrast, 19/BB cells demonstrated equivalent chromatin opening and transcription at *PDCD1* during activation but closed this site to pre-stimulation levels as they became dysfunctional (**Figure 2j**), suggesting divergent regulation of this exhaustion-defining site. Finally, we quantitated the similarity of these two cell states to a previously defined exhaustion-associated chromatin signature (*23*) and again found significantly greater overlap between open chromatin sites in classically exhausted cells and dysfunctional 19/28 (*P* = 8.82 × 10^−5^) and minimal overlap with dysfunctional 19/BB (*P =* 0.419, **Figure 2k**). Collectively, we observed that chronic stimulation of both CARs leads to T cell dysfunction but that chronic stimulation of 19/28 activates similar genomic programs to chronic TCR stimulation, while chronic stimulation of 19/BB leads to activation of a distinct regulatory program.

### Chronic stimulation of 19/BB promotes development of a unique dysfunctional cell state

To interrogate the transcriptional changes associated with 41BB-driven T cell dysfunction in a manner that would enable clarification of population heterogeneity, we performed longitudinal single cell RNA sequencing of chronically stimulated 19/28 and 19/BB. We observed that cells collected at days 0, 6 and 15 segregated into twelve clusters (**Figure 3a**) and that clusters were largely defined by timepoint of collection (**Figure 3b**). For example, clusters 2, 4, 5 and 6 were primarily resting cells, clusters 0, 1, 3, 7 and 10 were activated cells, while clusters 8 and 9 were dysfunctional cells. Pathway analysis using KEGG confirmed representation of predictable biologically pathways in each cluster (ie. cell cycle and metabolism genes in “activated” clusters 0, 1, 7 and 10, **Extended Data Fig. 3a**). Given our interest in understanding the transcriptional identities of dysfunctional CAR T cells, we focused on clusters 8 and 9. While cluster 9 contained roughly equivalent quantities of cells from both day 15 samples (44% 19/BB, 42% 19/28), cluster 8 was heavily skewed towards day 15 19/BB cells (83% 19/BB, 17% 19/28, **Figure 3b**). Evaluation of each sample individually demonstrated largely equivalent cluster distributions at days 0 and 6 (**Extended Data Fig. 3b**), but distinct cluster distribution at day 15 (**Figure 3c**), similar to our bulk sequencing analysis (**Figure 2a**). More than half of dysfunctional 19/28 occupied cluster 9, which was defined by high expression of exhaustion-associated genes. In contrast, ∼70% of dysfunctional 19/BB were in cluster 8, whose identity was defined by a distinct set of genes with varied roles in lymphocyte function including cytotoxicity (*GNLY, CCL5, PRF1, GZMA, GZMK, CTSW*), NK cell identity (*KLRK1, KLRC2*) and T cell differentiation (*ID2*). Expression of NK markers has previously been reported in dysfunctional endogenous (*24, 25*) and CAR T cells (*6*), although their contribution to dysfunction remains unclear. Consistent with the modest increase in classical exhaustion genes observed from bulk RNA sequencing (**Figure 2d**), we observed that a fraction of dysfunctional 19/BB occupied cluster 9, suggesting that dysfunctional 19/BB cells are heterogenous and contain some classically exhausted cells but that most have a novel transcriptional identity. Our bulk RNA sequencing and CyTOF data demonstrated that dysfunctional 19/BB cells expressed genes and proteins associated with T cell memory. We found that these genes were drivers of clusters 2 and 4 and, predictably, were enriched in day 0 cells (**Extended Data Fig. 4**). In day 15 samples, expression of these genes was found in cells that localized to cluster 8, although in a geographical subset of cluster 8 that overlapped with cluster 4 (**Extended Data Fig. 4**).

**Figure 3.**
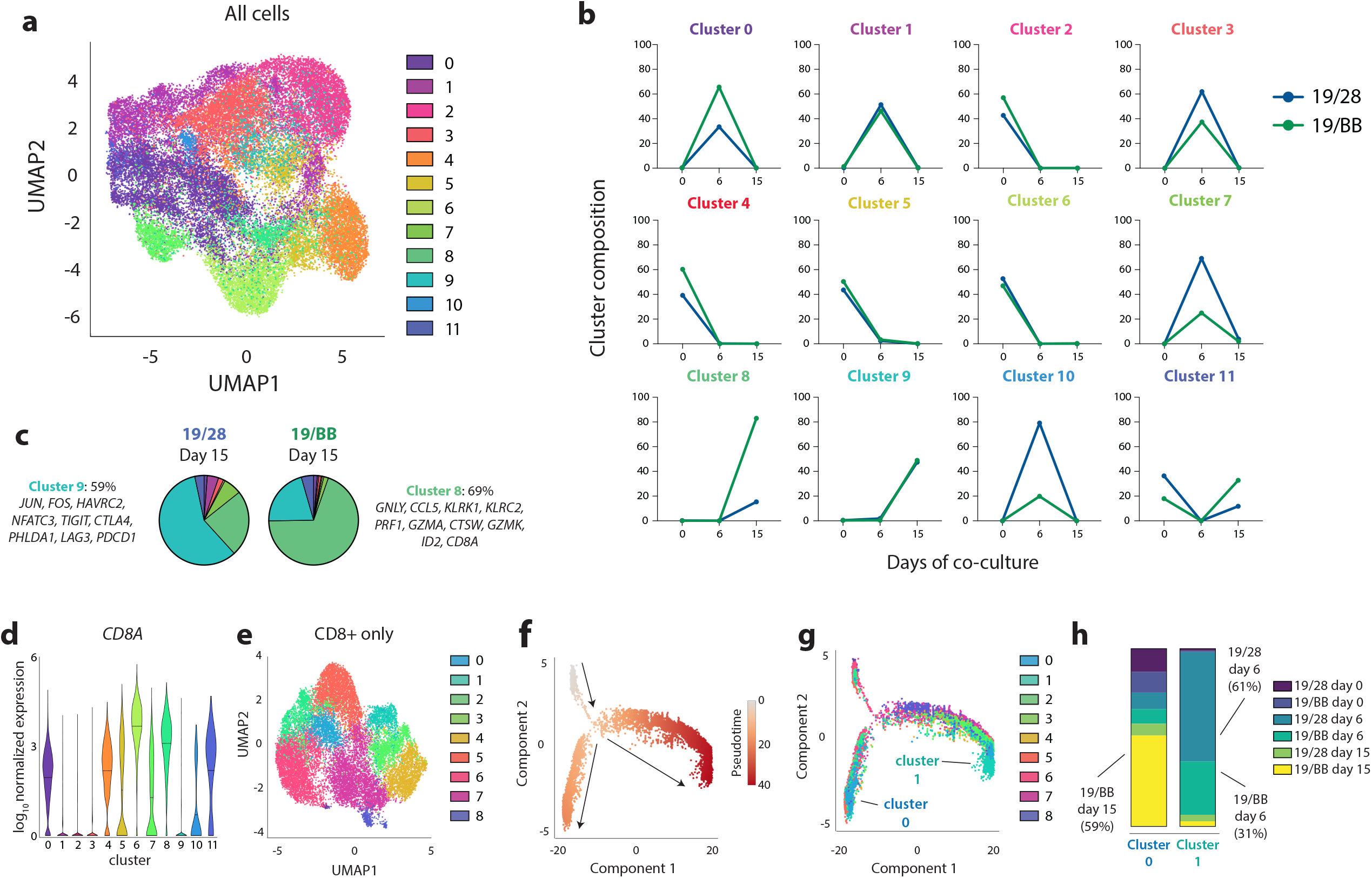
Single cell analysis reveals that dysfunctional 41BB CAR T cells are a unique terminal state. **a**, UMAP of 19/28 and 19/BB cells over time. **b**, Proportion of each sample (19/28 or 19/BB at day 0, 6 or 15) contained within each cluster. **c**, Proportion of each cluster in day 15 samples. **d**, *CD8A* expression in each cluster. **e**, Re-clustering of *CD8A*-expressing 19/28 and 19/BB cells from days 0, 6 and 15. **f**, Pseudotime analysis of CD8 cells. **g**, Mapping of CD8 clusters onto pseudotime. **h**, Proportion of each sample contained in terminal CD8 clusters 0 and 1.

As cluster 8 cells expressed uniformly high levels of *CD8A* (**Figure 3d**) we re-clustered only CD8 cells from all six samples. CD8s distributed to nine new clusters (**Figure 3e**) and we found that dysfunctional 19/28 cells primarily occupied CD8-cluster 7, defined by expression of *GZMB, IL2RA, ENO1, CCL3* and *BATF3* (**Extended Data Fig. 5a**). Dysfunctional 19/BB were highly enriched for CD8-cluster 0, defined by expression of *GNLY, KLRB1, CCL5, ID2* and *GZMK* (**Extended Data Fig. 5a**), reflecting similarity in identity of CD8-cluster 0 to cluster 8 from the whole population analysis (**Extended Data Fig. 5b**). To trace the transcriptional evolution of CD8+ CAR T cells as they progressed from resting to dysfunctional, we performed pseudotime analysis using Monocle2 (*26*). This demonstrated a divergence along two developmental trajectories (**Figure 3f**). Mapping of the CD8 clusters onto pseudotime revealed two terminal states at the conclusion of these trajectories, primarily composed of either CD8-cluster 0 or CD8-cluster 1 (**Figure 3g**). CD8-cluster 0 was highly-enriched for day 15 19/BB cells (59%, **Figure 3h**), confirming that this terminal state represented, in large part, dysfunctional 19/BB cells. While CD8-cluster 0 did contain some day 15 19/28 (6.8% of cluster total), these cells were distributed throughout the other clusters, primarily occupying CD8-cluster 6 (32% of day 15 19/28 total), which localized to the lower pseudotime trajectory but was pre-terminal (**Figure 3g**). CD8-cluster 1, in contrast, was highly similar to cluster 7 from the whole population analysis, both of which were defined by expression of activation markers *MKI67, BHLHE40* and *TOP2A* (**Extended Data Fig. 5c**). Consistent with these observations, CD8-cluster 1 was almost entirely (92%) cells from day 6 (**Figure 3h**). These findings suggest that chronic stimulation of 19/BB directs a distinct terminal T cell identity.

### CAR T cells that fail clinically acquire a dysfunctional 19/BB gene signature

Using our bulk RNAseq and scRNAseq datasets, we generated a gene signature of dysfunctional 19/BB cells (T_BBD_, **Extended Data Fig. 6, Supplementary Table 3**). To determine if this transcriptional state developed in CAR T cell products that failed to induce remission in patients, we analyzed circulating CAR T cells collected from a patient with diffuse large B cell lymphoma who received tisagenlecleucel, a commercially-available anti-CD19 CAR T cell product that contains the 41BB costimulatory domain. This patient had a partial response one month after treatment but progressive disease at three-month evaluation, a common clinical scenario that leads to persistent CAR stimulation. We evaluated peripheral blood cells circulating at the time of partial anti-tumor activity (day 14) and after therapeutic failure (day 100, **Figure 4a**). scRNAseq of CAR+ T cells revealed that day 14 cells clustered separately from cells collected at day 100, identifying a transition in transcriptional identity (**Figure 4b**). While the day 14 sample contained both CD4 and CD8 cells, the day 100 sample was composed almost entirely of CD8s (**Figure 4c**). Consistent with this observed loss of CD4s in failing cells, a recent report demonstrated that long-term response after 41BB-based CAR T cell treatment was predominantly driven by persistence of CD4+ CAR T cells (*27*). We assessed the overlap between the T_BBD_ signature and the genes that defined the identity of day 100 CAR T cells (upregulated compared to day 14, n=232) and observed significant overlap between these two gene sets (*P =* 2.8×10^−47^, **Figure 4d**). Further, we found that almost all of the top 10 drivers of the T_BBD_ signature were expressed at higher levels in day 100 cells than day 14 cells, with the exception of cytotoxicity genes *GNLY* and *PRF1* (**Figure 4e**). To assess if these cells also acquired features of exhaustion, we profiled expression of the master exhaustion genes defined from TIL datasets (*17-20*). We found that while 12 of these 17 genes were higher in day 100 cells, these differences were modest and none met a threshold log_2_-fold change >1.5 from day 14 to day 100 (**Figure 4f**). These observations mirror the modest increases in exhaustion-associated gene expression that arose in our *in vitro* model (**Figure 2d**). Overlap between Day 100 marker genes and TIL exhaustion signatures (*17-20*) revealed significantly less transcriptional similarity than observed for the T_BBD_ signature (**Figure 4g**). This suggests that failing 41BB-based CAR T cells evolve a transcriptional profile that more closely resembles the T_BBD_ gene profile more than an exhausted gene profile.

**Figure 4.**
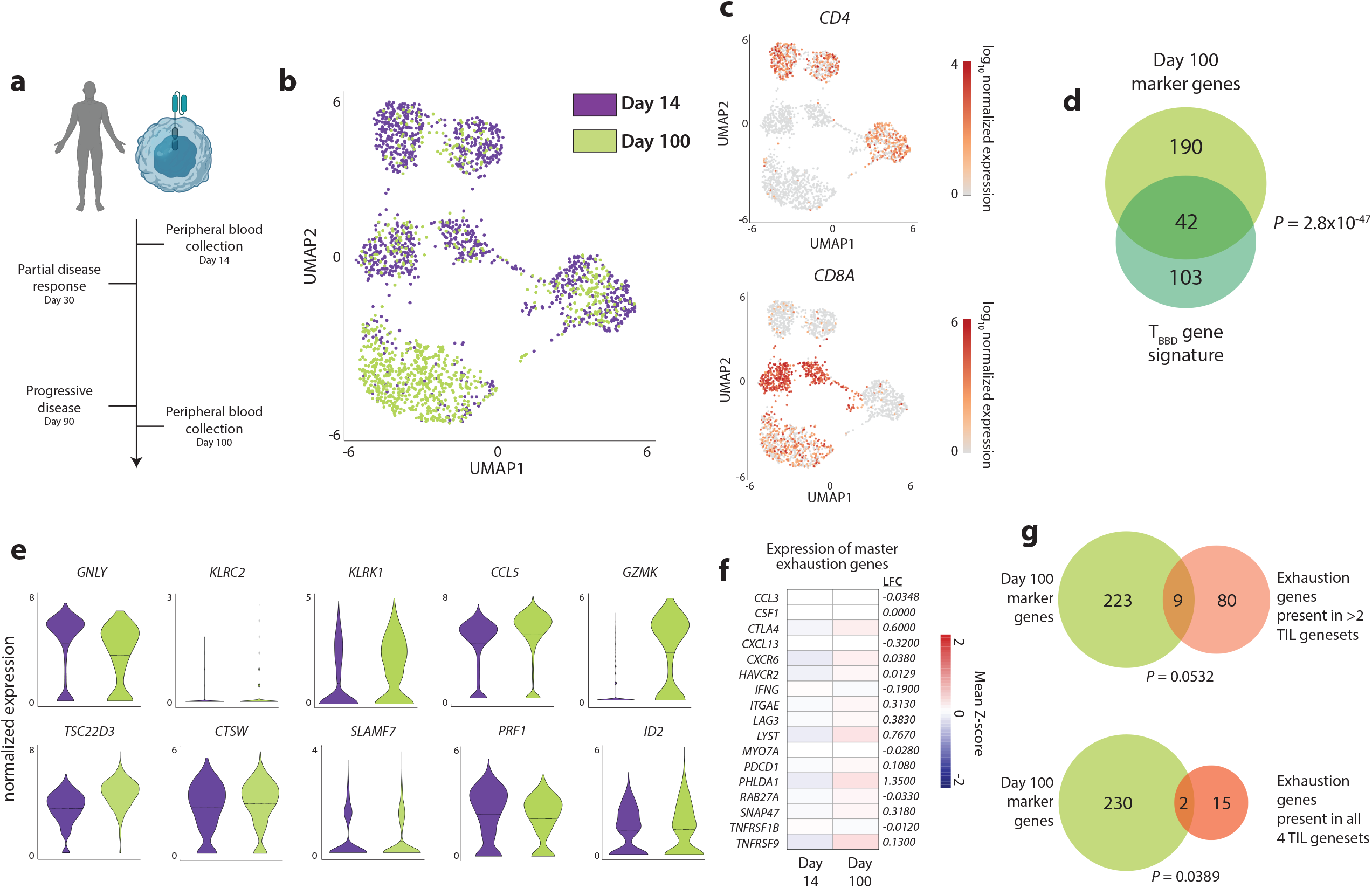
Failing CAR T cells express a dysfunctional 41BB CAR T cell signature. **a**, Peripheral blood cells from a patient who received tisagenlecleucel for diffuse large B cell lymphoma were collected 14 and 100 days after infusion and purified for CAR-expressing cells. **b**, UMAP of day 14 and day 100 cells. **c**, Expression of *CD4* and *CD8A* in peripheral blood CAR T cells. **d**, Overlap between *in vitro*-defined signature of 41BB-driven dysfunction and genes defining day 100 cell identity. Significance of overlap determined using Fisher’s Exact Test. **e**, Expression of Top 10 driver genes in day 14 and day 100 cells. **f**, Log_2_ fold change in expression of master exhaustion genes in day 100 compared to day 14 cells. **g**, Overlap between genes defining day 100 cell identity and TIL exhaustion signatures. Significance of overlap determined using Fisher’s Exact Test.

### Dysfunctional 19/BB cells re-activate FOXO3

To identify the biological pathways responsible for promoting the development of this transcriptionally unique state we performed transcription factor motif analysis using our ATACseq data. We specifically evaluated how binding site accessibility evolved as each cell type (19/28 or 19/BB) progressed from resting to dysfunctional. We found many of the top 100 transcription factor binding motifs with increased accessibility in dysfunctional cells were shared between 19/28 and 19/BB cells (32/100), but a greater number were specific to each cell type (68/100). As has been demonstrated previously (*8*), 19/28 cells opened sites for binding of the heterodimeric transcription factor AP1 (**Figure 5a**). 19/BB cells, in contrast, opened sites for binding of homeobox (HOX) and forkhead box (FOX) transcription factors (**Figure 5b**). Pathway analysis of the unique binding sites observed for each group confirmed that dysfunctional 19/28 cells were enriched for binding of basic leucine zipper domain (bZIP)-containing factors (which includes AP1 and BATF family members) as well as T-box factors, such as TBX21 (T-bet) (**Figure 5c**). Dysfunctional 19/BB cells were broadly enriched for HOX and FOX factor sites (**Figure 5d**). Interrogation of our RNAseq data revealed nearly no expression of any *HOX* genes in day 15 cells, but we did observe a resurgence in *FOXO* (**Figure 5e**) and some *FOXP* (**Extended Data Fig. 7a**) factors at day 15, which was more significant for 19/BB cells. We also found higher expression of bZIP factors in dysfunctional 19/28 cells, with previously observed increases in expression for *JUNB, FOS* and *FOSB* (*8*) (**Extended Data Fig. 7b**).

**Figure 5.**
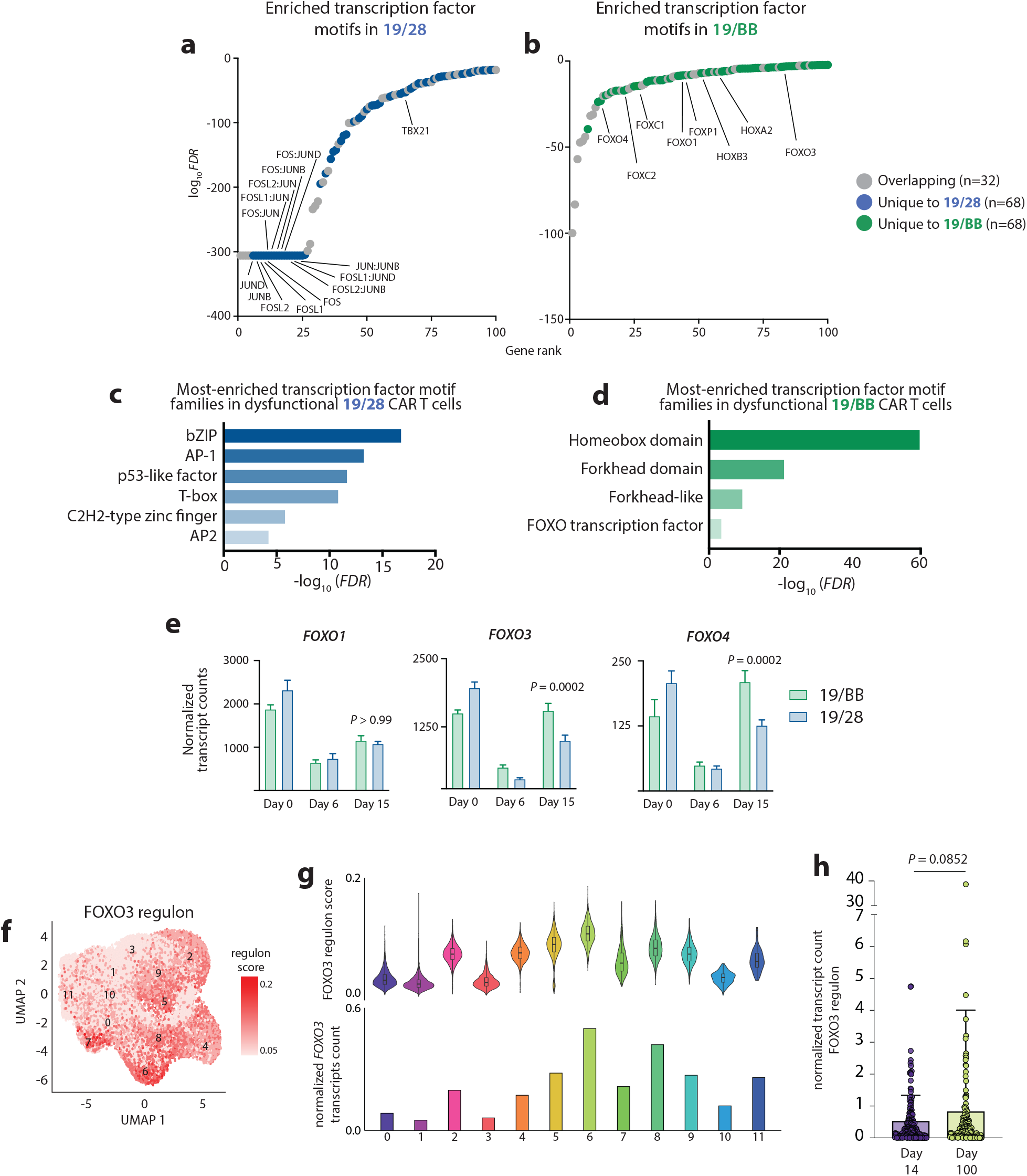
T_BBD_ cells demonstrate reactivation of FOXO3. **a-b**, Transcription factor motif analysis demonstrates increased accessibility of **a**, AP1 sites in 19/28 cells and **b**, HOX and FOX sites in 19/BB cells as cells progress from resting to dysfunctional. **c-d**, Pathway enrichment analysis of unique sites with increased accessibility in **c**, 19/28 and **d**, 19/BB cells. **e**, Expression of FOXO1, FOXO3 and FOXO4 transcripts over time. Significance determined using two-way ANOVA. **f**, Expression of the FOXO3 regulon in 19/28 and 19/BB cells collected at days 0, 6 and 15. **g**, FOXO3 regulon score for each scRNAseq cluster (above) and normalized *FOXO3* transcript count for each cluster (below). **h**, Expression of FOXO3 regulon in day 14 and day 100 cells collected from patient peripheral blood. Significance determined using Mann-Whitney Test.

We next sought to determine if the increased FOXO binding site accessibility and increased *FOXO* transcript quantity was accompanied by increased FOXO activity. To do this we used SCENIC, a computational tool designed to discriminate cell states based on expression of transcription factor gene regulatory networks using single cell data (*28*). This analysis mapped high activity of the FOXO3 regulon (target network) in clusters composed of resting cells (2, 4, 5 and 6), consistent with its known role in maintaining T cell homeostasis (*29*), as well as clusters 7 and 8 (**Figure 5f**). Quantification of regulon scores, reflecting quantity of FOXO3 target gene expression, from each cluster as well as quantification of average *FOXO3* transcript counts from cells in each cluster confirmed that both were highest in clusters 6 and 8, suggestive of high FOXO3 activity specifically in resting cells and dysfunctional 19/BB cells (**Figure 5g**). GSEA of cluster 8 marker genes further confirmed enrichment of the FOXO3 regulon (NES=2.71, *FDR* 0, **Extended Data Fig. 7c**). A key target of FOXO3 is *FOXP3* (*30, 31*), responsible for directing development of regulatory T cells (T_reg_). In addition to higher expression of CD25, a marker of T_regs_, in dysfunctional 19/BB (**Figure 1i**) we also observed higher expression of *FOXP3* transcripts in these cells (**Extended Data Fig. 7d**). Evaluation of our single cell data revealed that while *FOXP3* transcripts could be detected in cells from several clusters, they were most enriched in cluster 8 (**Extended Data Fig. 7f**), collectively validating the observed increase in FOXO3 activity using both transcript and protein expression of a known FOXO3 target. We evaluated FOXO3 target gene expression in peripheral blood CAR+ T cells and observed a trend towards increased FOXO3 activity from day 14 to day 100, however this was not statistically significant (*P =* 0.0852, **Figure 5h**). These data demonstrate that as 19/BB cells become dysfunctional they re-activate the transcription factor FOXO3.

### Disruption of FOXO3 improves function of chronically stimulated 19/BB cells

Among other functions, FOXO3 maintains T cell homeostasis and is rapidly inhibited upon T cell activation (*29*). To identify if FOXO3 was a direct inhibitor of 19/BB function, we disrupted genomic *FOXO3* in 19/28 or 19/BB using Cas9-based engineering (**Figure 6a**). This resulted in >90% gene disruption (**Extended Data Fig. 8a**) which did not impact T cell expansion during manufacturing (**Extended Data Figs. 8b-c**), consistent with previous studies of *Foxo3*^KO^ in murine T cells (*32*). We subjected these cells to our chronic *in vitro* stimulation assays and found that loss of FOXO3 modestly impaired expansion of 19/28 but not 19/BB (**Extended Data Fig. 8d-e**). Evaluation of anti-tumor cytotoxic function demonstrated that loss of FOXO3 did not alter potency or durability of 19/28 but improved function of 19/BB (**Figure 6b)**, delaying the onset of CAR T cell dysfunction in these short chronic stimulation cultures by several days (**Figure 6c**). To further validate the role of FOXO3 in dysfunction, we designed constructs that encoded either 19/28 or 19/BB CARs and a transgenic *FOXO3* (**Figure 6d**). Expression of these constructs resulted in overexpression of FOXO3 in CAR+ T cells (**Extended Data Fig. 8f**), and resulted in a minor improvement in cell expansion during manufacturing for both 19/28 and 19/BB (**Extended Data Fig. 8g**). In our chronic stimulation assays we observed that FOXO3 overexpression had minimal impact on 19/28 expansion, however it dramatically inhibited 19/BB expansion (peak CAR T cell expansion from baseline for 19/BB: 22.8x; for 19/BB *FOXO3*^OE^: 3.2x; *P < 0*.*0001*, **Extended Data Fig. 8h**). This impaired expansion was accompanied by a rapid and robust loss of tumor control for 19/BB cells (**Figure 6e**) and a much earlier onset of dysfunction (**Figure 6f**). Interestingly, we observed that *FOXO3*^OE^ cells were significantly smaller in size throughout chronic stimulation (**Extended Data Fig. 8j**), suggestive of a less-activated cell state.

**Figure 6.**
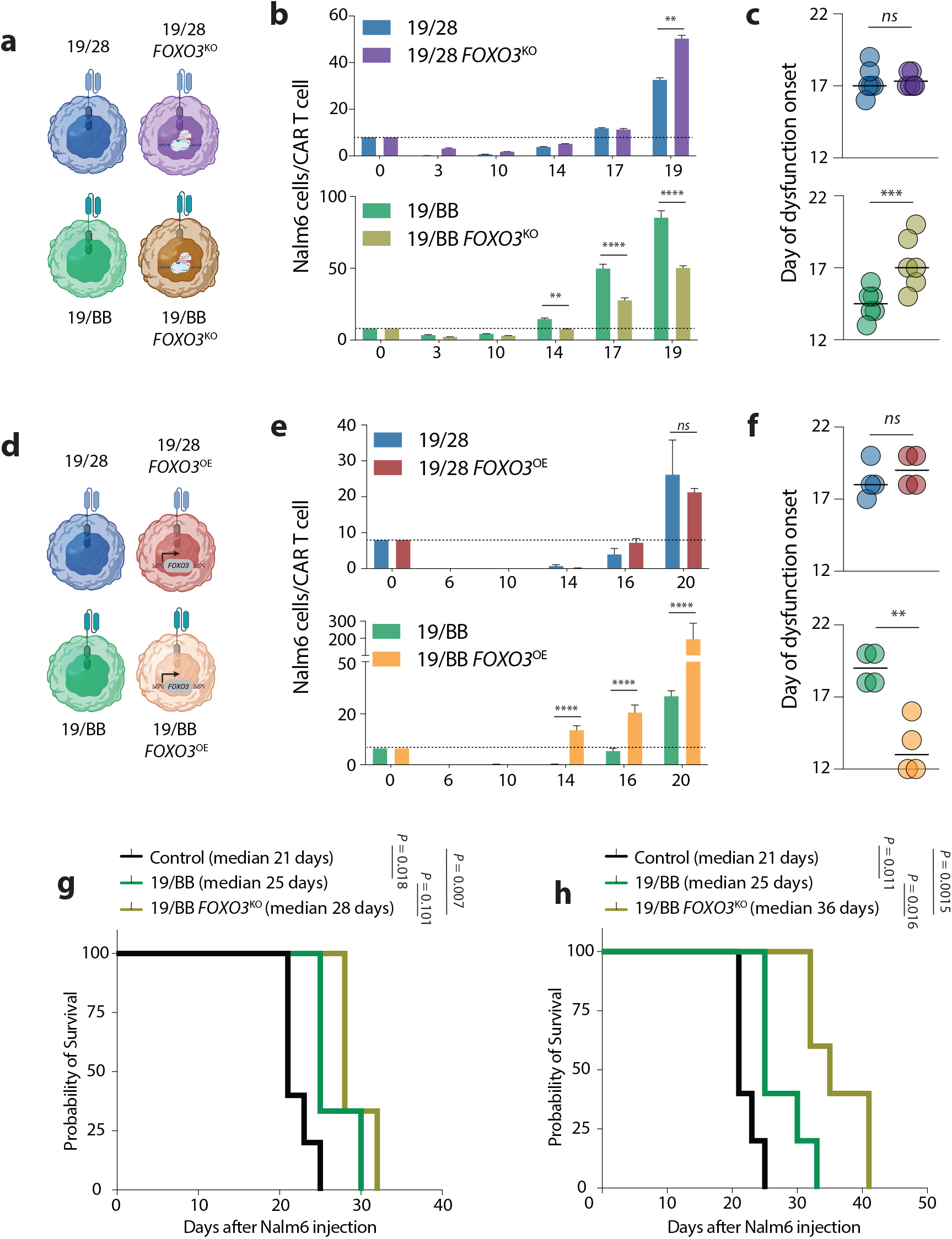
Manipulation of *FOXO3* impacts 41BB-based CAR T cell function. **a**, Representation of *FOXO3*^KO^ cells being evaluated. **b**, target Nalm6 cells per CAR T cells over the course of chronic stimulation. Representative data from n=3 donors. **c**, Day of dysfunction onset as measured by first day of T cell contraction or loss of tumor control. Data from n=3 independent donors. **d**, Representation of *FOXO3* over-expression cells being evaluated. **e**, target Nalm6 cells per CAR T cells over the course of chronic stimulation. Representative data from n=2 donors. **f**, Day of dysfunction onset as measured by first day of T cell contraction or loss of tumor control. Data from n=2 independent donors. **g**, Survival of mice after treatment with 0.125×10^6^ CAR T cells. Significance determined using Log-Rank test. **h**, Survival of mice after treatment with 0.5×10^6^ CAR T cells. Significance determined using Log-Rank test. For both experiments n=5 mice per group.

We proceeded to evaluate the impact of *FOXO3*^KO^ on 19/BB cells *in vivo*. We used two stress models of systemic human ALL (*33*), in which large tumor burdens are established but low T cell quantities are given, in order to recreate the high tumor burdens that have associated with clinical CAR T cell failure (*3*). NOD/SCID/γ^-/-^ (NSG) mice were engrafted with 10^6^ Nalm6 cells and then given sub-therapeutic doses of CAR T cells seven days later. In the first model we delivered 1.25×10^5^ CAR T cells and found that loss of FOXO3 has a modest impact in improving anti-leukemic activity (**Extended Data Fig. 9a**). This translated to a trend towards improved survival for mice who received *FOXO3*^KO^ cells (*P =* 0.101, **Figure 6g**). In a second stress model we delivered the same Nalm6 dose but gave 5×10^5^ CAR T cells on day seven. Here we observed greater anti-leukemic activity in both 19/BB groups, with significantly enhanced anti-tumor function of *FOXO3*^KO^ cells (**Extended Data Fig. 9b**). At this higher dose, *FOXO3*^KO^ 19/BB cells enabled a more impactful prolongation of animal survival (improvement in median survival from 25 days to 36 days with *FOXO3*^KO^, *P =* 0.0162, **Figure 6h**). Collectively, these data indicate that FOXO3 plays a key suppressive role in the anti-leukemic function of CAR T cells that is dependent on 41BB activation.

## Discussion

Chimeric antigen receptors have improved outcomes for many patients with refractory hematologic cancers, yet most patients are not cured by this therapy. Understanding how these synthetic proteins direct effective and defective T cell functionality is critical to broadening efficacy of this platform both for B cell cancers and other malignancies. Several studies have identified similarities between dysfunctional CAR T cells and exhausted T cells, however few have been designed to understand how CARs may direct divergent pathways of differentiation. Here, we demonstrate that CARs bearing the 41BB costimulatory domain direct the development of a unique state of T cell dysfunction that is, in part, reliant on FOXO3 activity. CAR-driven T cells differ from TCR-driven T cells in several critical ways. Most notably, chronically activated endogenous T cells do not undergo prolonged 41BB costimulation, as this receptor is only transiently expressed after initial activation. Thus, the persistent and high-intensity 41BB activation that arises in the setting of chronic CAR T cell stimulation may synthetically force circuitry that does not occur naturally. Given that four of six FDA-approved CAR T cell products are 41BB-based, identification of the mechanistic regulators of this novel biology is of significant relevance.

A notable phenotypic attribute of dysfunctional 19/BB cells is expression of genes encoding NK receptors, a finding that has been observed in various models of TCR-driven T cell exhaustion (*24, 25*). Interestingly, two recent reports identified a novel late-stage but pre-terminal exhausted state (termed ‘T_KLR_’) in the murine chronic LCMV model system (*34, 35*). These cells were defined by expression not only of NK surface receptors but also *Gzma Gzmk, Ccl5* and *Id2*, as well as lack of expression of terminal exhaustion markers *Pdcd1, Lag3* and *Tox*, reflecting a highly similar transcriptional identity to the T_BBD_ signature defined here. If T_KLR_ and dysfunctional 19/BB are a similar intermediate exhausted lineage is not clear – dysfunctional 19/BB notably lack expression of *CX3CR1*, found to be a defining marker of T_KLR_ cells (and all other intermediate exhausted states). Further, whether development of T_KLR_ cells is dependent on 41BB, as it is for CARs, or instead relies on a shared pathway independent of 41BB is the focus of ongoing studies. Intriguingly, both studies found that T_KLR_ cells were a fraction of the bulk exhausted T cell population, while we observe the majority of cells present at day 15 of chronic 19/BB stimulation bear the T_BBD_ gene signature. If indeed these cell states are dependent on 41BB signaling, this higher frequency may reflect the increased strength and duration of 41BB activation from CARs as opposed to natural costimulatory signals resulting from chronic infection.

A recent study similarly interrogated the trajectory of chronically stimulated 41BB-based CAR T cells (*6*). While the authors also observed expression of NK receptors in dysfunctional 41BB-based cells, they did not identify the same transcriptional features or regulatory pathways our studies did. Whether this is a result of a context dependence to this dysfunctional circuitry (their CARs targeted low-abundance mesothelin on the surface of pancreatic cancer cells in contrast to our model which targeted high-abundance CD19 on leukemia cells) remains unknown. The authors also observed increased expression of classical exhaustion markers in dysfunctional as compared to resting CAR T cells, as have others (*36, 37*). We, too, observe an increase in expression of these markers as 41BB-based CAR T cells progress from rest to dysfunctional, but identify that this expression is modest and restricted to a distinct, smaller subset of dysfunctional cells. These data highlight that low-level expression of exhaustion markers may not be the same as commitment to the classical exhaustion program; indeed, we found dysfunctional 19/BB to have closed chromatin at the *PDCD1* locus which would preclude defining these cells as classically exhausted. Broadly, phenotypic markers of any cell state should be considered in context. “PD1+” should likely be broken into a spectrum of “PD1^hi^” to “PD1^lo^”. While this can be difficult with clinical tissues, increased precision will enhance our ability to distinguish between an increasingly complex spectrum of T cell lineages. Notably, we do not observe expression of NK receptor genes in CD28-based CAR T cells, suggesting this is not a broad feature of CAR-driven dysfunction but is dependent on costimulatory structure.

Forkhead box, also known as winged helix, transcription factors have broad roles in an array of cellular functions across tissue types, including T cells. By inducing *FOXP3* expression, FOXO3 promotes differentiation of induced T_reg_ (*30, 31*). Our data demonstrate an enrichment of CD25+ and FOXP3+ cells in dysfunctional 19/BB. Two recent reports identified that enrichment of T_reg_ cells is associated with clinical failure of CAR T cells, directly implicating a role for this pathway in limiting anti-tumor efficacy. These clinical data reflected evaluation of pre-infusion and early post-infusion CD28-based anti-CD19 CAR T cells, thus the contribution of induced T_reg_ on the impaired functionality of 19/BB remains to be determined. Beyond T_reg_ development, FOXO3 has been shown to suppress anti-viral T cell function by inhibiting the formation of durable memory in patients with HIV (*38*) and in murine models of LCMV (*39*). Of particular relevance to the findings presented here, a distinct report identified that deletion of *Foxo3* in T cells significantly improved T cell control of chronic LCMV (*40*). The authors found that loss of Foxo3 had no impact on effector function early during infection and also did not impact T cell expansion through chronic stimulation, both of which we observed as well. These data strongly bolster our findings by corroborating a role for FOXO3 in a distinct model of chronic antigen receptor stimulation. How FOXO3 exerts this suppressive function remains unclear. Several candidate pathways exist, including promoting apoptosis, promoting a resting-like homeostatic state, or altering T cell metabolism (*29*). Of specific interest to the development of next-generation CAR therapies is the link between 41BB and FOXO3. We were surprised to observe that manipulation of FOXO3 did not impact T cell manufacturing or the function of CD28-based CARs, both of which are driven by CD3/CD28 signaling. These findings support the hypothesis that FOXO3’s suppressive effect is dependent on 41BB signaling and underscore that elucidating the relationship between 41BB, FOXO3 and activation of the T_BBD_ dysfunction program is of critical importance.

We were intrigued by the observation that FOXO3 loss did not improve 19/BB expansion but did improve anti-tumor cytotoxicity, while, in contrast, FOXO3 overexpression impacted both expansion and cytotoxicity. Understanding the biology responsible for this nuanced difference is the focus of ongoing studies but these data highlight that manipulation of transcription factors is far from precise. These proteins operate in complex regulatory networks, and we speculate that compensatory regulatory pathways may restrict 41BB-based CAR T cell expansion in the setting of FOXO3 loss. Regardless, these data demonstrate that disrupting *FOXO3* does not result in unconstrained 19/BB expansion but improved anti-tumor function on a per-CAR T cell basis, findings with significant translational relevance.

While beneficial, disruption of *FOXO3* resulted in modest delay in the onset of 41BB-based CAR T cell failure in our *in vitro* and *in vivo* stress models. This is consistent with the improvements seen in other studies that disrupt T cell transcription factors in an effort to improve CAR efficacy (*6*), and suggests that an individual factor is unlikely to completely rescue CAR-driven T cell dysfunction. Multiplexed manipulations that exploit vulnerabilities in distinct pathways of CAR-driven circuitry are likely necessary to propel synergistic improvements in function. Another limitation of this study is the inclusion of clinical samples from only one patient. Given the nature of our question, the role of chronic CAR stimulation on T cell dysfunction, we prioritized sequencing of cells undergoing chronic stimulation for a prolonged period. Most studies evaluate CAR T cells collected between days 7-30 after infusion (*41, 42*), or cells collected at later time points from patients with durable responses, a clinical setting in which CAR T cells are not chronically stimulated and persist longer (*37, 43*). To our knowledge, this is the first study to report data on CAR T cells isolated months after infusion from a patient with progressive lymphoma, a scenario in which CAR T cell persistence is highly limited beyond 30 days. While we demonstrate feasibility of evaluating these late-time point cells, this analysis required a significant volume of peripheral blood which is not available for many patients enrolled on current banking protocols. Given the important biology we were able to identify using these cells, we aim to improve collection procedures moving forward to enhance our ability to evaluate these valuable tissues.

## Acknowledgements

The authors wish to thank Abby Green, Julie Ritchey, Matthew Cooper, Alun Carter, John DiPersio, Hamza Celik, Grant Challen, Nicole Helton, Tim Ley and Carl DeSelm for helpful discussions and technical assistance. This work was supported by P50CA171963 and R01CA205239 (T.A.F.), K08CA237740, Be The Match Amy Strelzer Manasevit Research Award, Gabrielle’s Angel Foundation Cancer Research Award and Damon Runyon Clinical Investigator Award (all to N.S.).

## Author contributions

M.E.S., J.H.L. and N.S. designed and planned experiments. M.E.S, J.H.L, J.L., A.H., Y.-S.H, T.-C.C., J.C., J.W., H.H. and N.S. performed experiments. N.K., G.H., M.M.B.-E., M.F., S.K.-S., D.D. and T.A.F. contributed to CyTOF studies. M.S. coordinated primary sample acquisition and analysis. M.E.S. and A.G. performed bulk RNA and ATAC sequencing analysis. M.T. and M.A. performed single cell sequencing analysis. M.E.S. and N.S. wrote the manuscript. All authors reviewed the manuscript.

## Data availability

All bulk RNA, bulk ATAC and single cell RNA sequencing data from pre-clinical studies will be available through GEO (accession number forthcoming). Requests for clinical sample sequencing data will be reviewed by Washington University to determine if they are subject to confidentiality obligations. All other data generated from this study will be made available upon request to the corresponding author.

## Figure Legends

**Extended Data Figure 1.**
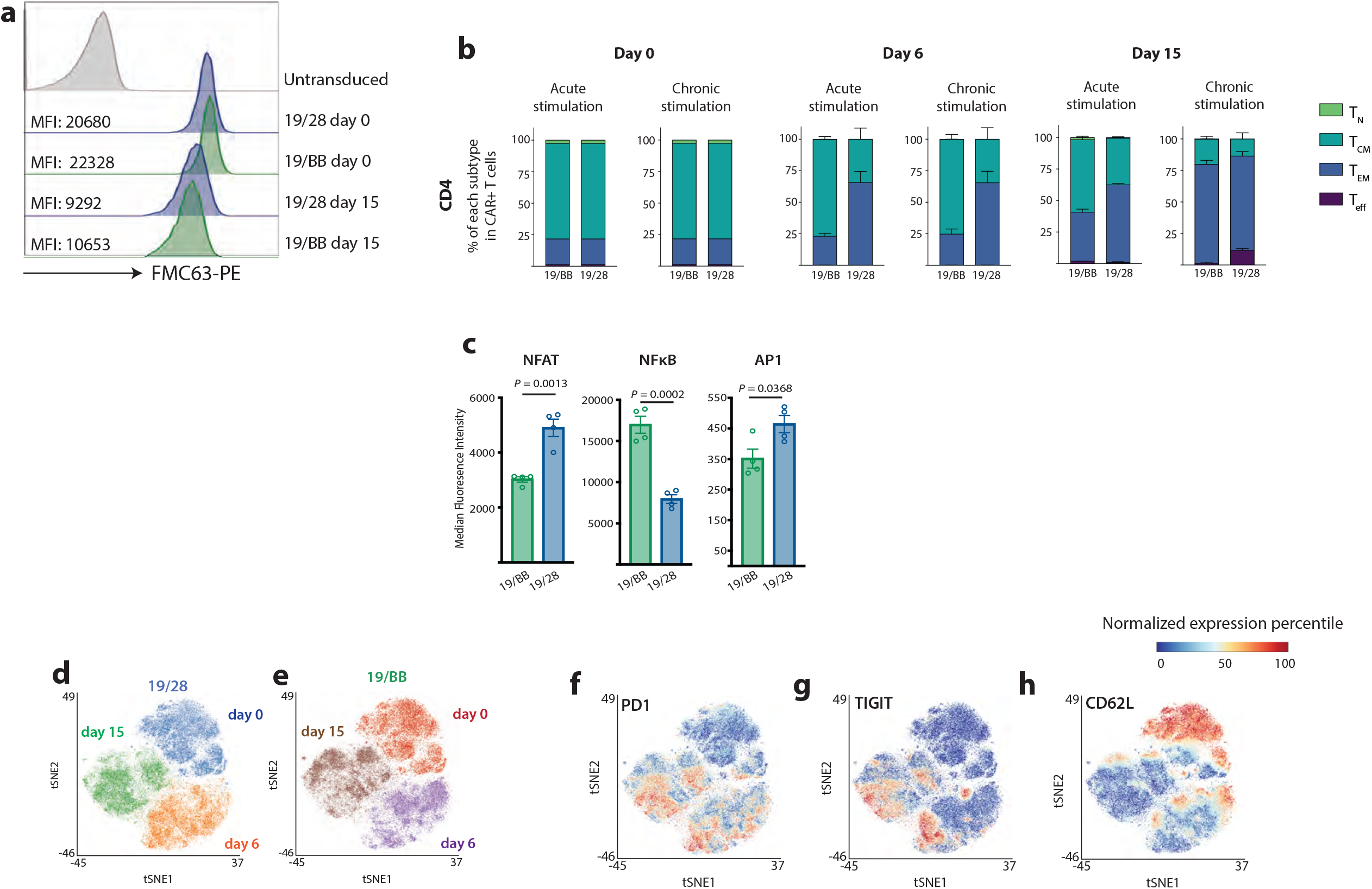
**a**, CAR expression in 19/28 and 19/BB after purification before co-culture (day 0) and after 15 days of chronic stimulation (day 15). **b**, Change in memory phenotype of CD4+ CAR T cell products after either acute (single combination with Nalm6 cells) or chronic stimulation. **c**, Activation of central T cell transcription factors in CAR Jurkat cells engineered to express a triple fluorescent reporter system. **d-e**, tSNE projection of **d**, 19/28 and **e**, 19/BB cells evaluated by CyTOF. Expression of **f**, PD1, **g**, TIGIT and **h**, CD62L in 19/28 and 19/BB cells.

**Extended Data Figure 2.**
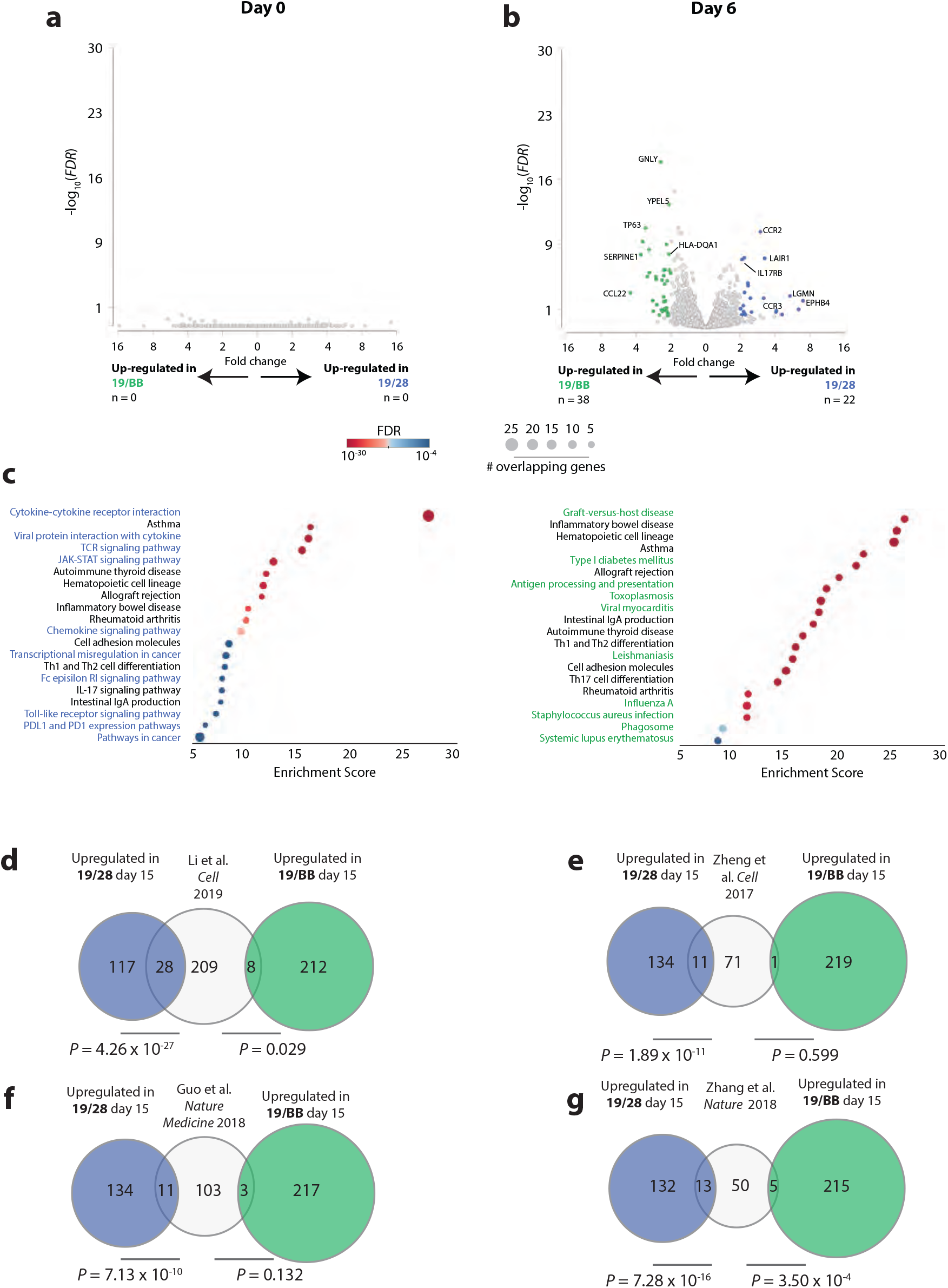
**a-b**, Volcano plot of DEGs at **a**, day 0 and **b**, day 6. **c**, KEGG pathways enriched in dysfunctional 19/28 and 19/BB cells. **d-g**, Overlap of genes with higher expression in 19/28 and 19/BB with previously published genesets defining exhausted tumor-infiltrating lymphocytes. Significance determined using Fisher’s Exact Test.

**Extended Data Figure 3.**
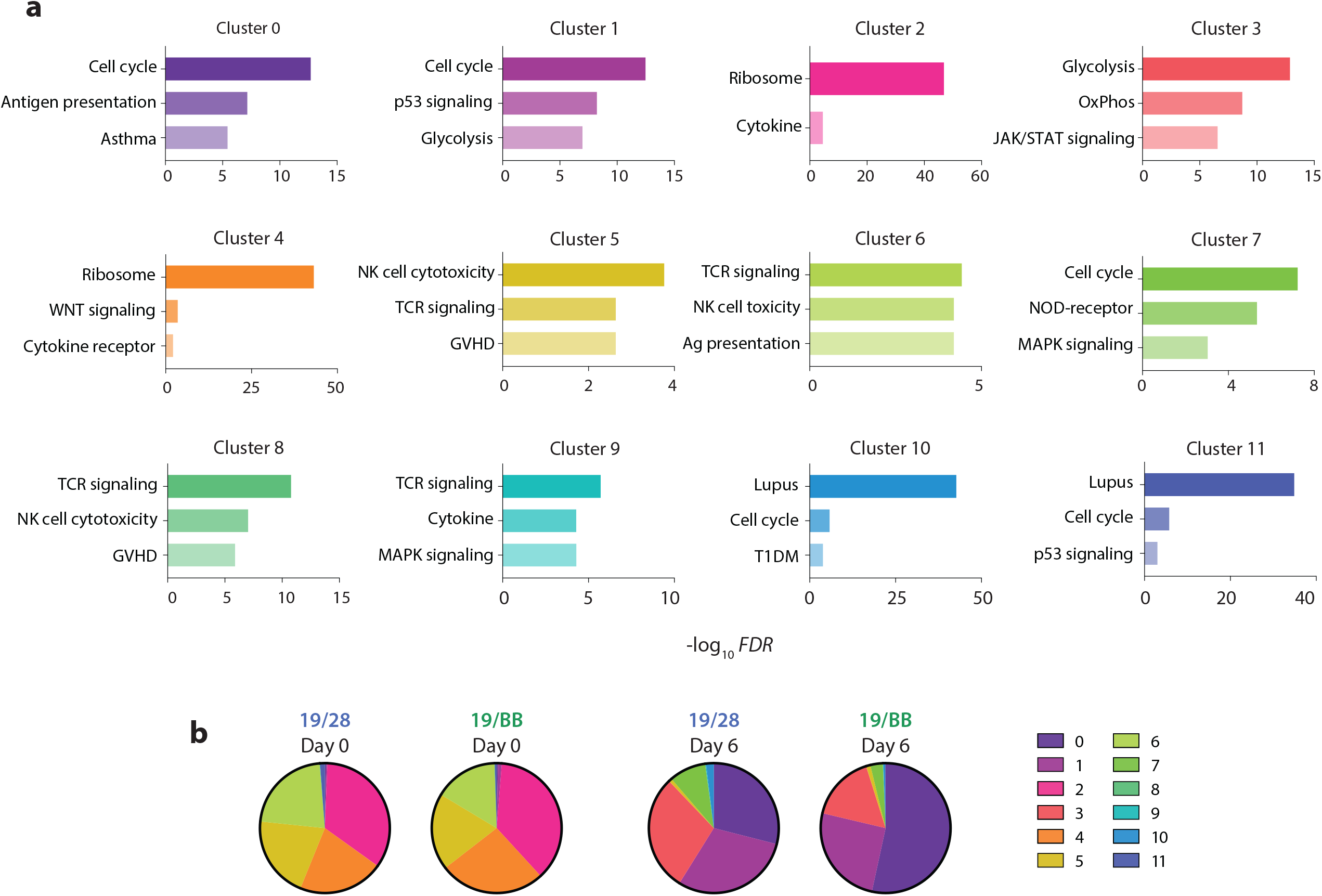
**a**, KEGG pathways enriched in each cluster. **b**, Proportion of each cluster present in day 0 and day 6 samples.

**Extended Data Figure 4.**
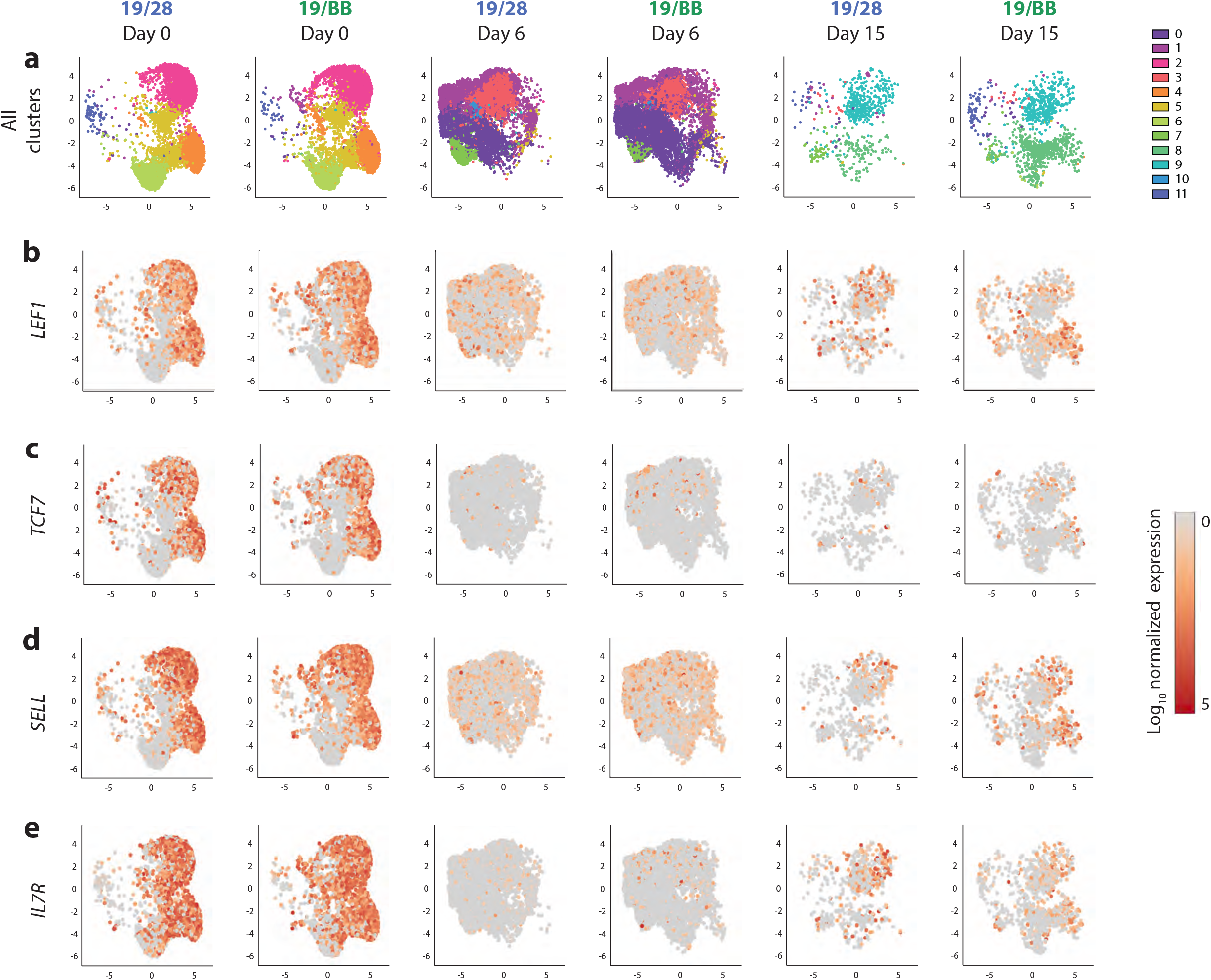
**a**, UMAP of scRNAseq for each individual sample. Expression of memory genes **b**, *LEF1*, **c**, *TCF7*, **d**, *SELL* and **e**, *IL7R*.

**Extended Data Figure 5.**
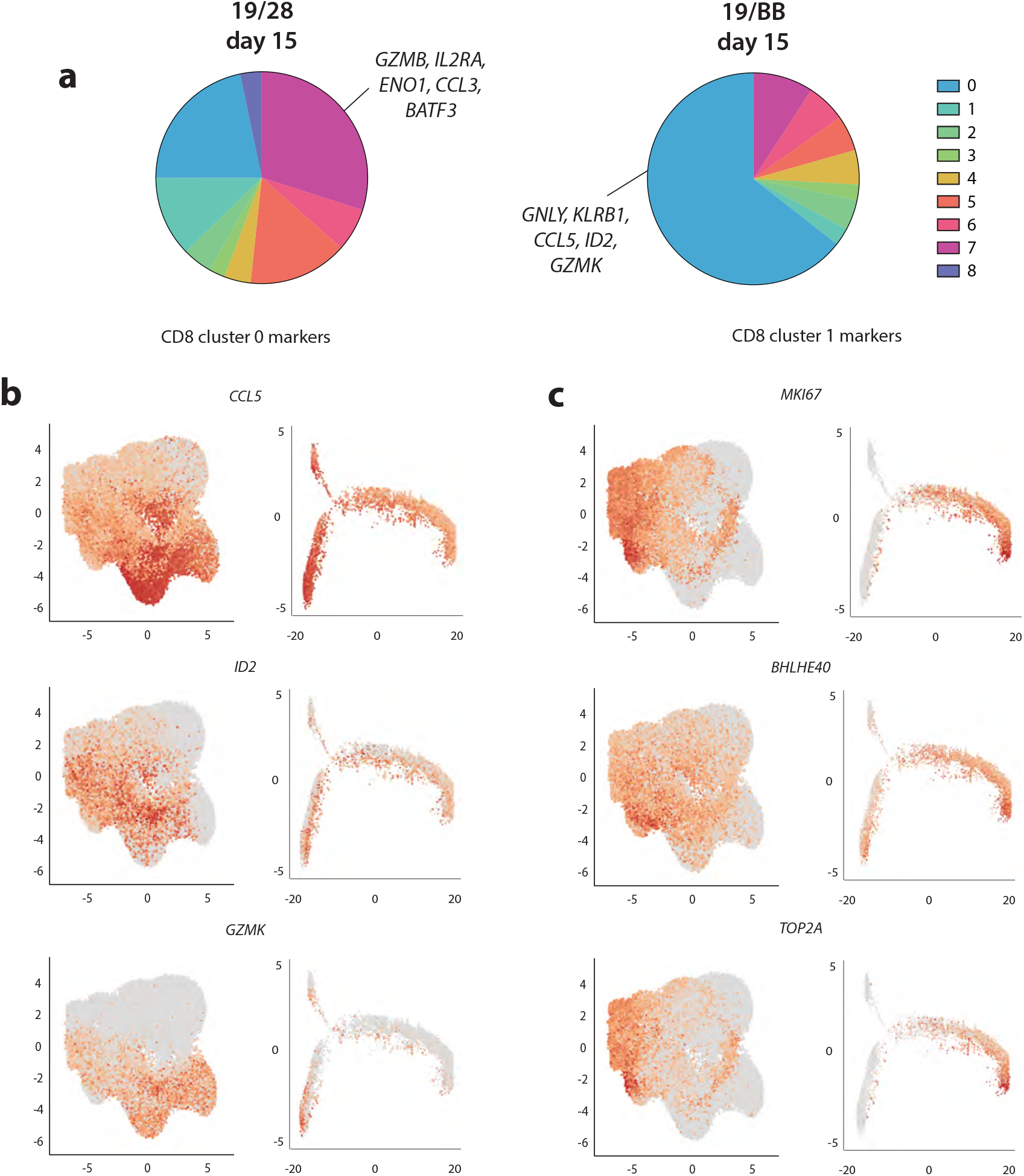
**a**, Proportion of each CD8 cluster in day 15 samples. **b-c**, Expression of CD8 **b**, cluster 0 and **c**, cluster 1 markers in whole population analysis and in pseudotime analysis of CD8 cells.

**Extended Data Figure 6.**
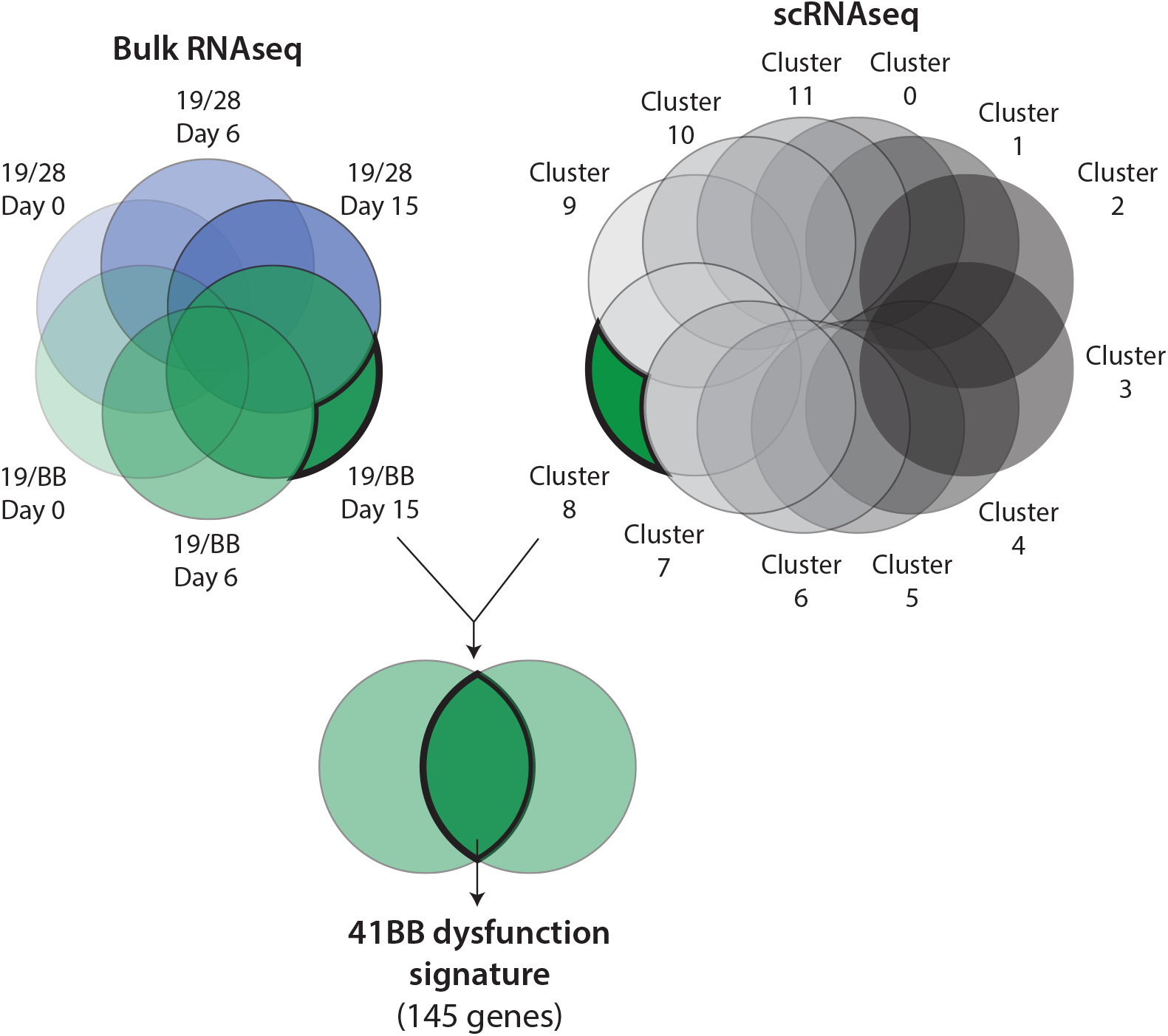
Approach to generate a signature of 41BB-driven CAR T cell dysfunction. Genes that were uniquely upregulated in day 15 (dysfunctional) 19/BB cells by bulk RNAseq were compared to genes that were uniquely upregulated in cluster 8 from our scRNAseq dataset. Genes that were shared from these two lists were used to generate the 41BB dysfunction signature of 145 genes. Filtering to identify genes in both datasets was performed with *FDR* < 0.05 and log_2_-fold change >1.5.

**Extended Data Figure 7.**
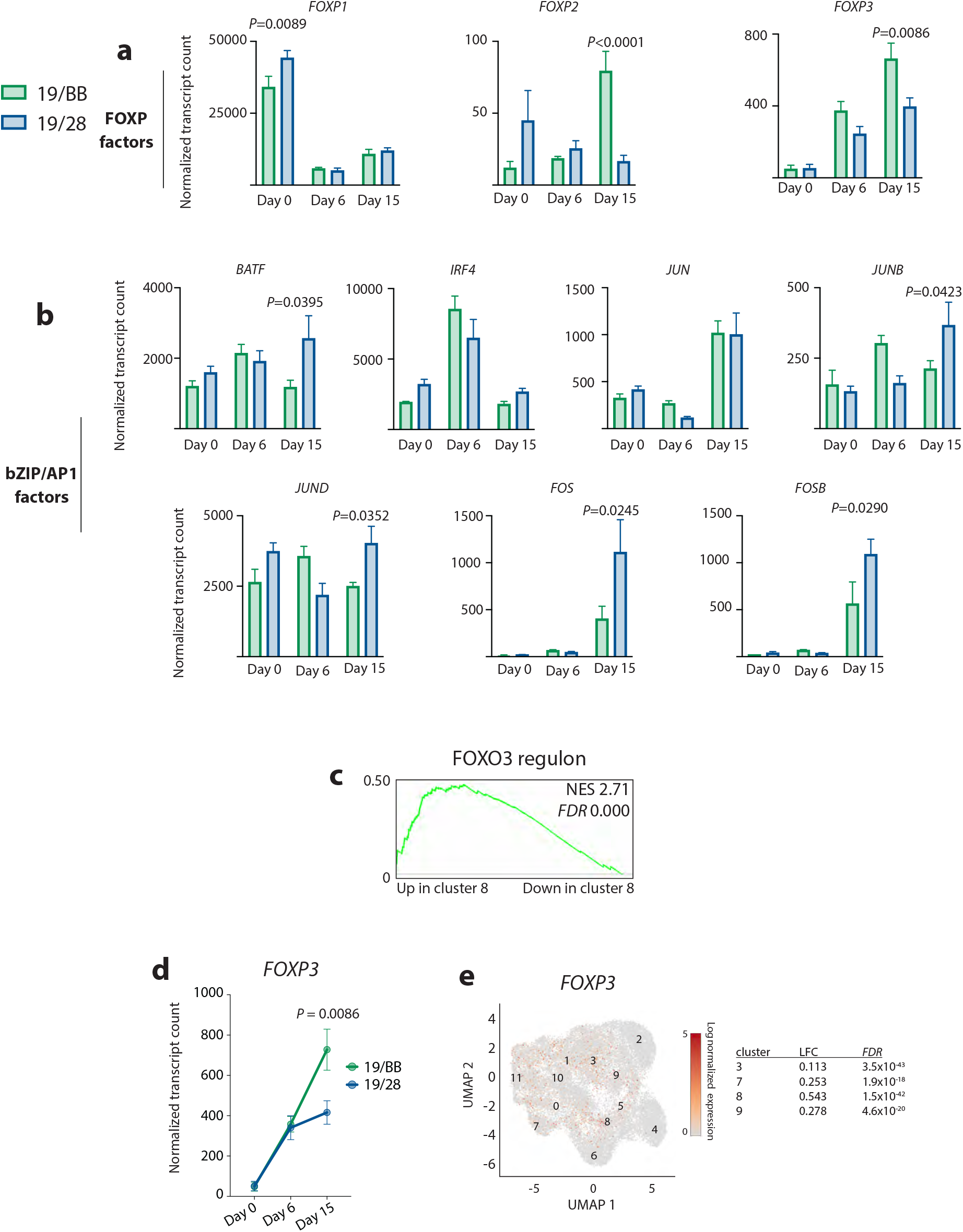
**a**, Expression of *FOXP* transcripts over time. **b**, Expression of bZIP/AP1 factors over time. **c**, Enrichment of FOXO3 target genes in genes that marked identity of cluster 8. **d**, Expression of *FOXP3* transcripts over time in chronically stimulated 19/28 and 19/BB cells. **e**, Expression of *FOXP3* in scRNAseq of 19/28 and 19/BB cells collected at day 0, 6 and 15. Table on left represents log_2_-fold change (LFC) in expression of FOXP3 in each cluster that it is found to be enriched with associated false discovery rate (FDR).

**Extended Data Figure 8.**
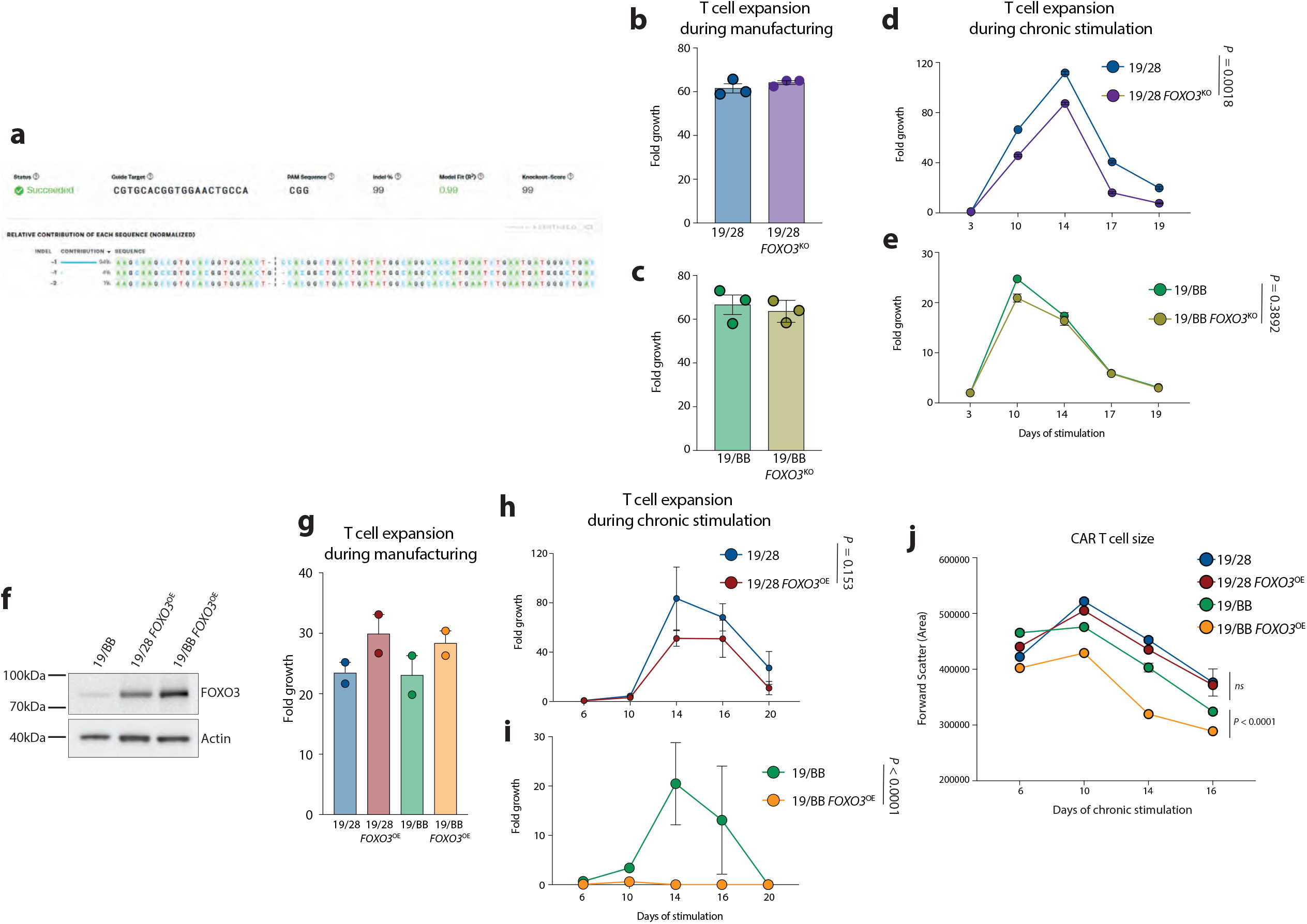
**a**, Sequencing analysis of genomic *FOXO3* in CAR T cell manufacturing products demonstrating high-efficiency knockout. Performed using Synthego ICE. **b**, T cell expansion during manufacturing of 19/28 or **c**, 19/BB cells with genomic disruption of *FOXO3*. N=3 independent donors. **d**, Expansion of *FOXO3*^KO^ or WT 19/28 or **e**, 19/BB CAR T cells during chronic stimulation. Representative data from n=3 independent donors. **f**, Western blot of lysates from CAR T cells engineered to overexpress FOXO3. **g**, T cell expansion during manufacturing of 19/28 and 19/BB with overexpression of FOXO3. N=2 independent donors. **h**, Expansion of *FOXO3*^OE^ 19/28 or **i**, 19/BB CAR T cells during chronic stimulation. Representative data from n=2 independent donors. **j**, CAR T cell size as measured by forward scatter area over the course of chronic stimulation. Representative data from n=2 independent donors. Statistical significance determined by two-tailed ANOVA with Bonferroni correction for multiple comparisons.

**Extended Data Figure 9.**
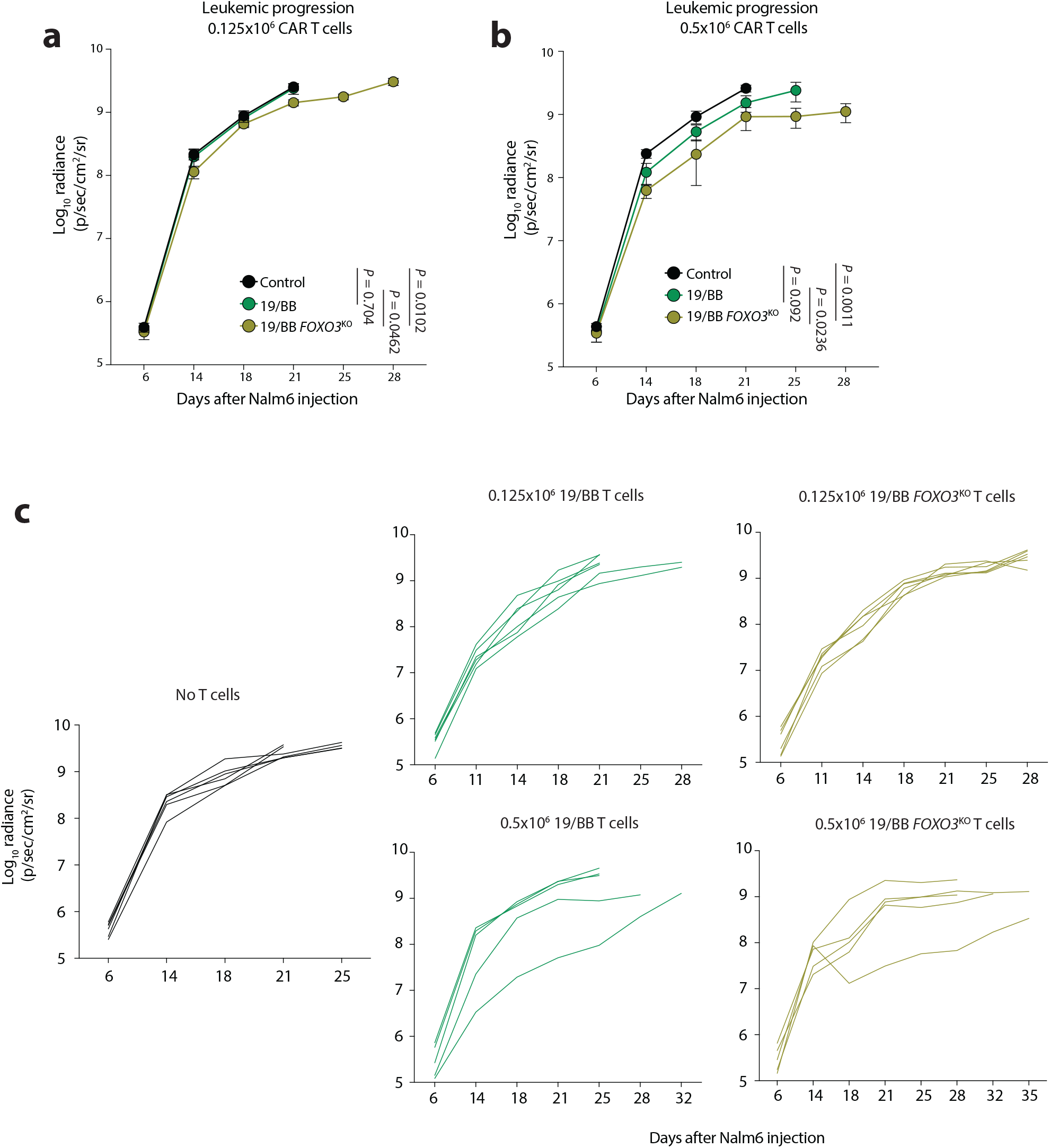
**a**, Nalm6 progression over time after treatment with 0.125×10^6^ CAR T cells or **b**, 0.5×10^6^ CAR T cells. Radiance curves were stopped at time of first animal death. Significance determined using two-way ANOVA. **c**, Individual animal radiance over time. For all studies n=5 animals per group.

## Methods

### CAR T cell manufacturing and chronic stimulation cultures

Lentiviral vectors were manufactured as previously described (*9*). PBMCs were procured from Miltenyi Biotec and CD4 and CD8 cells were purified using magnetic beads (Miltenyi Biotec) and combined at a 1:1 ratio. T cells were activated using CD3/CD28 stimulatory beads (DynaBeads; Thermo-Fisher) at a ratio of 3 beads/cell and incubated at 37°C overnight. The following day, CAR lentiviral vectors were added to stimulatory cultures at a MOI of 2-4. Beads were removed after 6 day of stimulation, and cells were counted daily until growth kinetics and cell size demonstrated they had rested from stimulation. For all studies described, CAR constructs also encoded a truncated CD34 (tCD34) surface marker of transduction, separated from the CAR transgene by a P2A sequence. Both CD28 and 41BB-based CARs were composed of the FMC63 single chain variable fragment targeting CD19, CD8α hinge and transmembrane regions, followed by the CD3ζ signaling domain and a terminal costimulatory domain.

Chronic stimulation cultures were established by combining CAR T cells (0.5-2×10^6^) with GFP+ Nalm6 cells at a 1:8 effector to target ratio in standard media. Cultures were profiled every other day by flow cytometry to both count cells and evaluate changes in protein expression. The co-cultures were re-fed with additional GFP+ Nalm6 cells every other day to maintain 1:8 effector to target ratio until the onset of dysfunction. CAR T cells were purified prior to co-culture at day 0, at day 6-7 when they are at the peak of activation and day 13-17 when they become dysfunctional for down-stream analyses.

### General cell culture and flow cytometry

Unless otherwise specified, cells were grown and cultured at a concentration of 1×10^6^ cells/mL of standard culture media (RPMI 1640 + 10% FCS, 1% penicillin/streptomycin, 1% HEPES, 1% non-essential amino acids) at 37°C in 5% ambient CO_2_. All co-culture studies were performed at an effector cell to target cell ratio of 1:8, unless otherwise stated. Samples were stained with CD34 (BD, clone 581, #555824), CD4 (Biolegend, clone OKT4, #317444), CD8 (Invitrogen, clone SK1, #17-0087-42), PD1 (Biolegend, clone eh12.2h7, #329928), TIM3 (Invitrogen, clone F38-2E2, #17-3109-42), LAG3 (Biolegend, clone 11c3c65, #369314), CD62L (Biolegend, clone dreg-56, #304822), CD45RO (Biolegend, clone UCHL1, #204236) and 7-AAD (BD, #559925) in 100ul FACS buffer (2%FBS in PBS), washed once with the same buffer and analyzed on the Attune NxT Flow Cytometer (ThermoFisher). GFP+ Nalm6 and GFP-CD34+ CAR T cells were gated and analyzed using FlowJo v9 or 10 (BD Biosciences).

### CyTOF

Mass cytometry was performed as previously described (*44*). Briefly, isolated CAR+ T cells were live/dead stained with a short pulse of cisplatin and surface stained for 30 minutes at room temperature. Cells were then washed and fixed overnight at 4°C with fix/perm buffer (eBiosciences). Intracellular staining was performed the following day at 4°C for 1 hour. Cells were barcoded according to manufacturer’s instructions (Fluidigm). Cells were washed and suspended in PBS containing 2% paraformaldehyde with Cell-ID Intercalator-IR. Mass cytometry data was collected on a Helios mass cytometer and analyzed using Cytobank (Beckman Coulter).

### Bulk RNA and ATAC sequencing and data analysis

Total RNA was extracted using Qiazol (Qiagen) and recovered by RNA Clean and Concentrator spin columns (Zymo). Samples were prepared according to library kit manufacturer’s protocol, indexed, pooled, and sequenced on an Illumina NovaSeq 6000. Basecalls and demultiplexing were performed with Illumina’s bcl2fastq2 software. RNA-seq reads were then aligned and quantitated to the Ensembl release 101 primary assembly with an Illumina DRAGEN Bio-IT on-premise server running version 3.9.3-8 software. All gene counts were then imported into the R/Bioconductor package EdgeR (*45*) and TMM normalization size factors were calculated to adjust for samples for differences in library size. The TMM size factors and the matrix of counts were then imported into the R/Bioconductor package Limma. Weighted likelihoods based on the observed mean-variance relationship of every gene and sample were then calculated for all samples and the count matrix was transformed to moderated log 2 counts-per-million with Limma’s voomWithQualityWeights. Differential expression analysis was then performed to analyze for differences between conditions and the results were filtered for only those genes with Benjamini-Hochberg false-discovery rate adjusted p-values less than or equal to 0.05. Further analysis was performed using Partek Flow (Partek Inc). Geneset Enrichment Analysis was done using GSEA v4.1.0.

Omni ATAC-seq libraries were made as previously described (*46*). Briefly, nuclei were isolated from 50,000 sorted CART19 cells, followed by the transposition reaction using Tn5 transposase (Illumina) for 30 minutes at 37°C with 1000rp mixing. Purification of transposed DNA was completed with DNA Clean and Concentrator (Zymo) and fragments were barcoded with ATAC-seq indices. Final libraries were double size selected using AMPure beads prior to sequencing. Paired-end sequencing (2 × 75 bp reads) was carried out on an Illumina NextSeq 500 platform. Adapters were trimmed using attack (version 0.1.5, https://atactk.readthedocs.io/en/latest/index.html), and raw reads were aligned to the GRCh37/hg19 genome using bowtie with the following flags: --chunkmbs 2000 --sam --best --strata -m1 -X 2000 (*47*). MACS2 was used for peak calling with an FDR cutoff of 0.05. Downstream analysis and visualization, including transcription factor motif analysis, was done using Partek Flow (Partek Inc).

### Single cell RNA sequencing

CAR T cells from chronic stimulation cultures were isolated using flow-based sorting as described. For clinical samples, frozen vials of peripheral blood from a patient who underwent CAR T cell therapy and experienced a transient partial response followed by disease progression were gently thawed, counted, and dead cells were removed (Dead Cells Removal kit, Miltenyi, #130-090-101). Resulting cell samples had viabilities of 92-98% and were stained using an anti-FMC63 antibody (Acro Biochemicals, clone Y45, #FM3-HPY53) and then enriched for CAR+ cells using flow-based sorting. These cells were then processed using the 10x Genomics Chromium Single Cell V(D)J Reagent Kits (10x Genomics, PN-1000006, PN-1000020, PN-120262) to generate single-cell emulsions for barcoding, reverse transcription and cDNA amplification. Immediately following these steps, 10x barcoded fragments were pooled and attached to standard Illumina adaptors to generate a barcoded single-cell RNA library. Sequencing libraries were quantified by qPCR before sequencing on the Illumina platform using HiSeq 4000 instrument.Cell Ranger’s count pipeline v6.1.1 (available at https://support.10xgenomics.com/single-cell-gene-expression/software/pipelines/latest/using/count) was applied to align reads and quantify gene expression of individual samples. Downstream single-cell analysis was performed using Seurat package v4.0.5 within the R programming environment v4.1.2 (*48*). Lower bound for the number of genes in individual cells was chosen based on binary logarithm distribution and was set to 9.8 for the dataset with chronic stimulation samples and 10.5 for the dataset with clinical samples. Additionally, cells with more than 7,500 genes for the dataset with chronic stimulation samples and 4,000 genes for the dataset with clinical samples were filtered out. The percentage of mitochondrial counts was calculated for every cell, and only cells with mitochondrial percentage less than 10% were used in further analysis. Filtered matrices were normalized using a scaling factor of 10,000 and centered. Two sources of unwanted variation, total number of counts and percentage of counts belonging to mitochondrial genes, were regressed using a linear model. Within the datasets samples were combined using the harmony batch correction function. UMAP dimensional reduction and shared nearest neighbor graph were calculated on harmony corrected PCA embeddings using 20 principal components. Number of principal components was selected based on the elbow plot. Graph-based clustering was performed on the reduced data. Differential expression analysis between clusters was performed using the MAST algorithm of Seurat R package (*49*), p value adjustment was done using Bonferroni correction. CD8+ T cells from the chronic stimulation dataset were processed with Monocle2 pseudotime analysis pipeline (*26*). Seurat object were converted into CellDataSet object and used as an input. Differentially expressed genes between the clusters were identified with generalized linear model MAST, filtered by a significance level of p adjusted < 0.05 and used for cell ordering. Dimensionality reduction was performed with DDRTree method. The cells from the day 0 were set as the root of the trajectory. Single-cell regulatory network analysis was performed with pySCENIC (*28, 50*). First, Seurat objects with raw filtered counts were converted into AnnData files via SaveH5Seurat and Convert functions from the SeuratDisk package (https://github.com/mojaveazure/seurat-disk/). Next, the adjacency matrix for transcription factors (hg38) and its targets were created using the GRNBoost2 algorithm (*51*). Motif analysis was performed using the cisTarget database (https://resources.aertslab.org/cistarget/). Cellular enrichment for each regulon was calculated by the AUCell module (*28*) with default thresholds. Visualization and downstream analysis of pySCENIC output were performed with the SCopeLoomR package (https://github.com/aertslab/SCopeLoomR/).

### T_BBD_ gene signature development

We determined which genes were biomarkers, or specifically enriched, in day 15 19/BB bulk RNA data and cluster 8 from scRNAseq data using Partek Flow. The overlap of each data set was used to generate the 145 gene signature. For these genes we developed a “contribution score” by multiplying the fold change of each gene relative to all other samples (5 samples from bulk RNAseq and 11 clusters for scRNAseq) from bulk and scRNAseq data. The 10 genes with the largest score were used for analysis in Figure 4e.

### CRISPR/Cas9 gene editing

CRISPR sgRNAs were designed using the CRISPick tool from The Broad Institute and the sgRNA design tool from Integrated DNA Technologies (IDT). Cells were electroporated using the Lonza 4D-Nucleofector Core/X Unit. Triple Reporter Jurkat cells were electroporated using the SE Cell Line 4-D Nucleofector Kit, and primary T cells were electroporated using the P3 Primary Cell 4-D Kit (Lonza). For Cas9 and sgRNA delivery, a ribonucleoprotein (RNP) complex was first formed by incubating 5µg of TrueCut Cas9 Protein V2 (Lonza) with 10µg of sgRNA for 10 min at room temperature. Cells were washed twice with PBS (Gibco) and spun at 100xg for 10 min and resuspended at a concentration of 2-10×10^6^ cells/100µL in the specified buffer. The RNP complex and 100µL of resuspended cells were combined and electroporated. Pulse codes EO-115 and CV-104 were used for resting primary T cells and Jurkat cells, respectively. After electroporation, the cells were incubated in standard 10% RPMI for Jurkat cells and cytokine enriched 10% RPMI (5 ng/ml IL7 and 5 ng/ml IL15, both from Peprotech) for primary T cells for the duration of experiment. Primary T cells were stimulated after 18 hours using CD3/CD28 Dynabeads (Thermo-Fisher) at 1:3 cell to bead ratio and engineered with lentivirus 24 hours later.

To determine efficiency of gene disruption, TIDE (Tracking of Indels by DEcomposition) analysis was used to detect knock out (KO) efficiency. Genomic DNA from electroporated cells was isolated (Qiagen DNeasy Blood & Tissue Kit) and 100-200ng were PCR amplified using Q5 Hot Start High Fidelity 2x Master Mix (NEB) and 10µM forward/reverse primers flanking the region of interest. Primers were designed such that the amplicon was at a target size ∼1 kb. PCR products were gel or column purified and sequenced, and trace files were analyzed using TIDE web tool (tide.nki.nl) to determine indel proportions. R^2^ values were calculated, reflecting goodness of fit after non-negative linear modeling by TIDE software (*52*).

### Xenograft mouse models

6-10 week old NOD-SCID-γc^-/-^ (NSG) mice were obtained from the Jackson Laboratory and maintained in pathogen-free conditions. Animals were injected via tail vein with 1×10^6^ Nalm6 cells in 0.2mL sterile PBS. On day 7 after tumor delivery, either 0.125×10^6^ or 0.5×10^6^ CAR+ T cells (WT or *FOXO3*^KO^) were injected via tail vein in 0.2mL sterile PBS. Animals were monitored for signs of disease progression and overt toxicity, such as xenogeneic graft-versus-host disease, as evidenced by >10% loss in body weight, loss of fur, diarrhea, conjunctivitis and disease-related hind limb paralysis. Disease burdens were monitored over time using the Xenogen IVIS bioluminescent imaging system, as previously described (*9*) and animals were sacrificed when radiance reached >3×10^9^ photos/sec/cm2/sr (5-log greater than background). To avoid skewing of radiance data, graphical representation for each group was stopped after death of the first animal in the group.

### Statistical analysis

All comparisons between two groups were performed using either a two-tailed unpaired Student’s t-test or Mann-Whitney test, depending on normality of distribution. Comparisons between more than two groups were performed by two-way analysis of variance (ANOVA) with Bonferroni correction for multiple comparisons. All results are represented as mean ± standard error of the mean (s.e.m.). Survival data were analyzed using the Log-Rank (Mantel-Cox) test.

---

**Supplementary Table 1:** CyTOF panel

| <u>Target</u> | <u>Tag</u> | <u>Clone</u> | <u>Manufacturer</u> | <u>Catalog #</u> |
| --- | --- | --- | --- | --- |
| CD45 | 089Y | HI30 | Fluidigm | 3089003B |
| Ki-67 | 115In | 8D5 | Thermo | MA5-15690 |
| CD3 | 141Pr | UCHT1 | Fluidigm | 3141019B |
| CD28 | 142Nd | CD28.2 | biolegend | 302902 |
| CD45Ra | 143Nd | H100 | Fluidigm | 3143006B |
| CD69 | 144Nd | FN50 | Fluidigm | 3144018B |
| CD4 | 145Nd | RPA-T4 | Fluidigm | 3145001B |
| CD8a | 146Nd | RPA-T8 | Fluidigm | 3146001B |
| RORgt | 147Sm | AFKJS-9 | ebioscience | 14-6988-82 |
| CD278/ICOS | 148Nd | C398.4A | Fluidigm | 3148019B |
| CD56 (NCAM) | 149Sm | NCAM16.2 | Fluidigm | 3155008B |
| CD223/LAG-3 | 150Nd | 11C3C65 | Fluidigm | 3150030B |
| CD103 | 151Eu | Ber-ACT8 | Fluidigm | 3151011B |
| TCRgd | 152Sm | 11F2 | Fluidigm | 3152008B |
| CD62L | 153Eu | DREG56 | Fluidigm | 3153004B |
| TIM3 | 154Sm | F38-2E2 | Fluidigm | 3154010B |
| CD27 | 155Gd | L128 | Fluidigm | 3158010B |
| CXCR3 | 156Gd | G025H7 | Fluidigm | 3156004B |
| CD134 (OX40) | 158Gd | ACT35 | Fluidigm | 3158012B |
| CD197 (CCR7) | 159Tb | G043H7 | Fluidigm | 3159003A |
| CD39 | 160Gd | A1 | Fluidigm | 3160004B |
| Tbet | 161Dy | 4B10 | Fluidigm | 3161014B |
| FoxP3 | 162Dy | PCH101 | Fluidigm | 3162011A |
| Eomes | 163Dy | WD1928 | ebioscience | 14-4877-82 |
| CD357 (GITR) | 164Dy | 621 | biolegend | 311602 |
| CD45RO | 165Ho | UCHL1 | Fluidigm | 3149001B |
| CD44 | 166Er | BJ18 | Fluidigm | 3166001B |
| Gata3 | 167Er | TWAJ | Fluidigm | 3167007A |
| CD154 (CD40L) | 168Er | 24-31 | Fluidigm | 3168006B |
| CD25 (IL-2R) | 169Tm | 2A3 | Fluidigm | 3169003B |
| CTLA-4 | 170Er | 14D3 | Fluidigm | 3170005B |
| Granzyme B | 171Yb | GB11 | Fluidigm | 3171002B |
| CD38 | 172Yb | HIT2 | Fluidigm | 3172007B |
| CD137/4-1BB | 173Yb | 4B4-1 | Fluidigm | 3173015B |
| CD279 (PD-1) | 174Yb | EH12.2H7 | Fluidigm | 3174020B |
| Perforin | 175Lu | B-D48 | Fluidigm | 3175004B |
| CD127 | 176Yb | A019D5 | Fluidigm | 3176004B |
| TIGIT | 209Bi | MBSA43 | Fluidigm | 3209013B |

**Supplementary Table 2:**
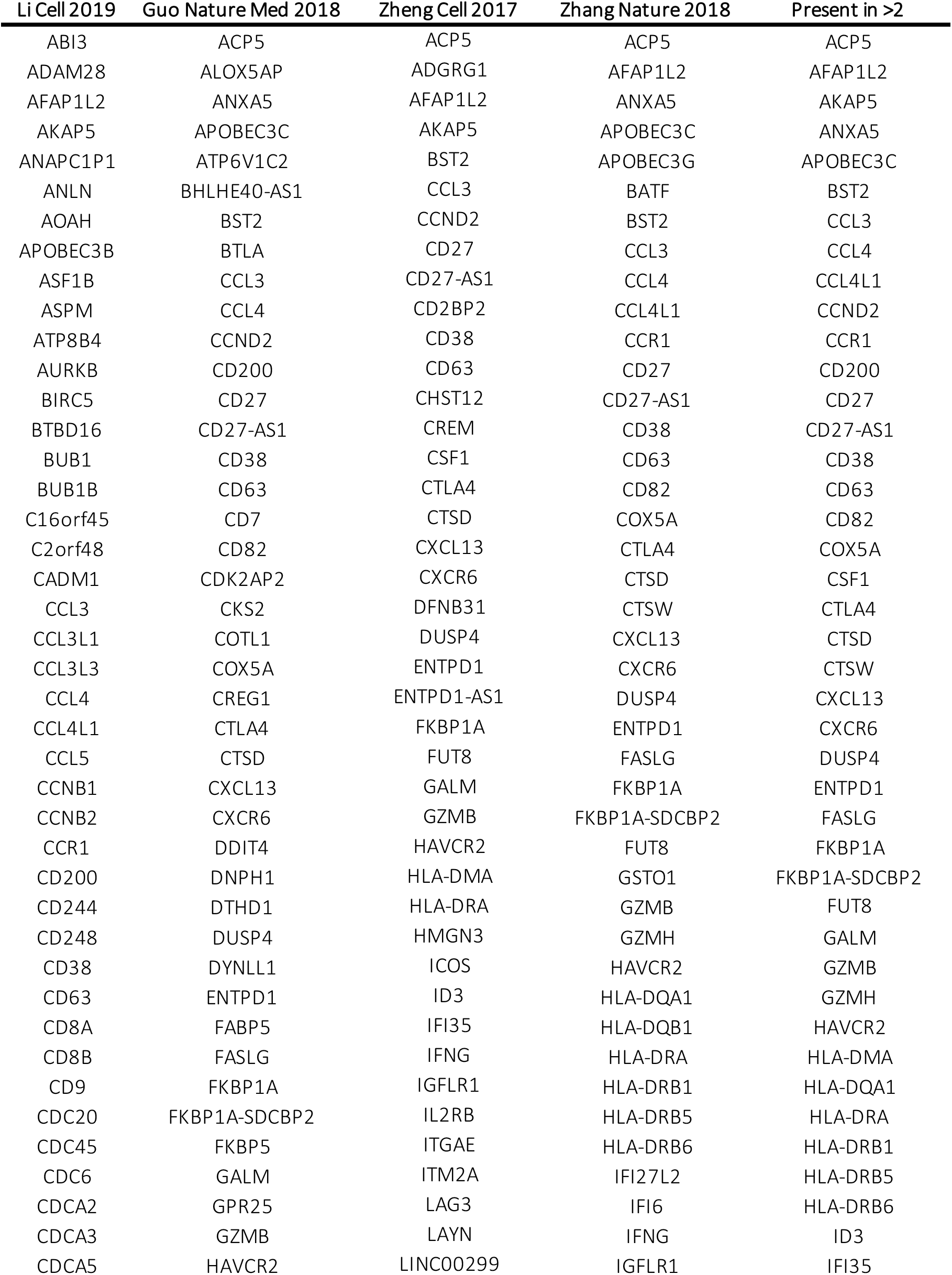
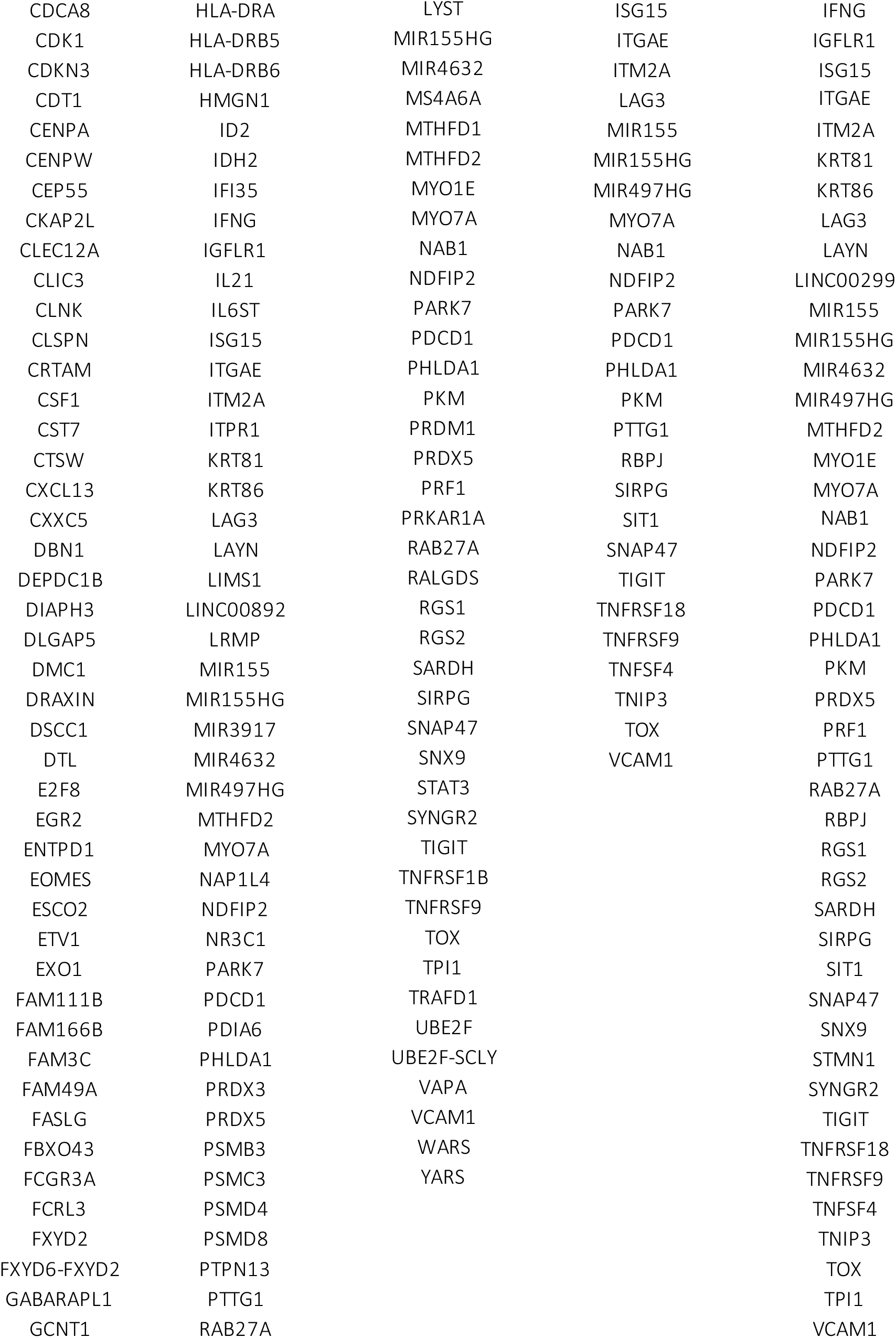
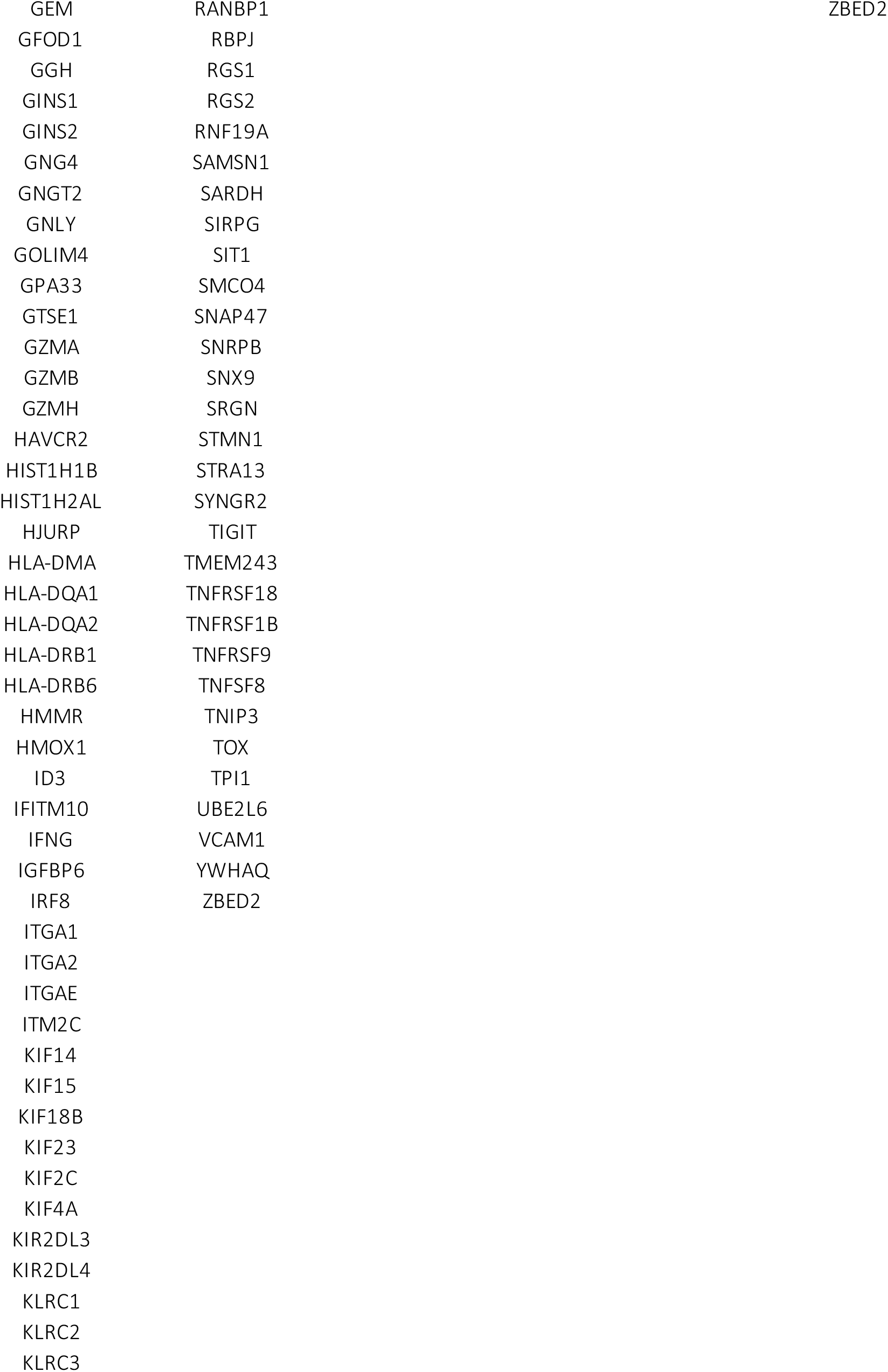
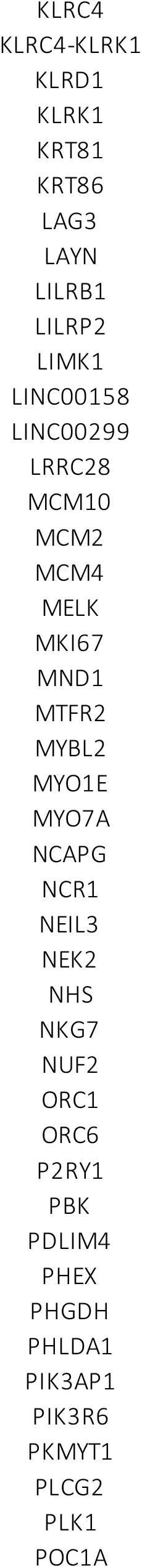
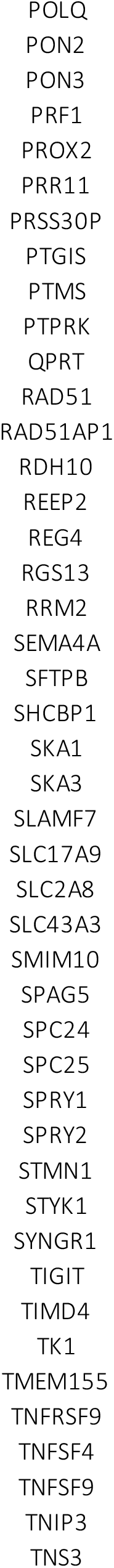
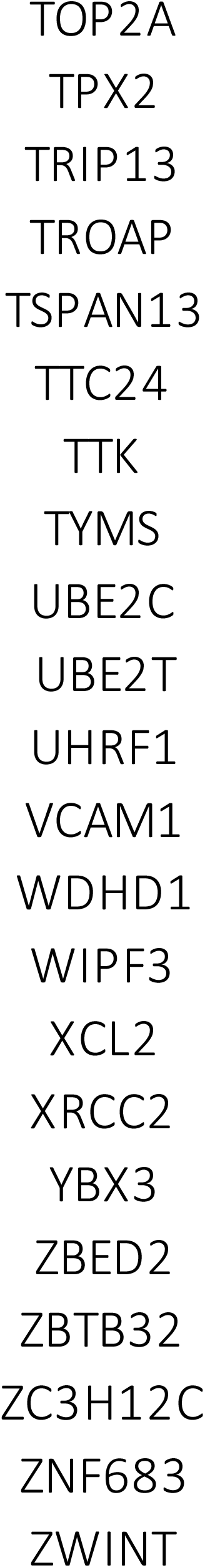
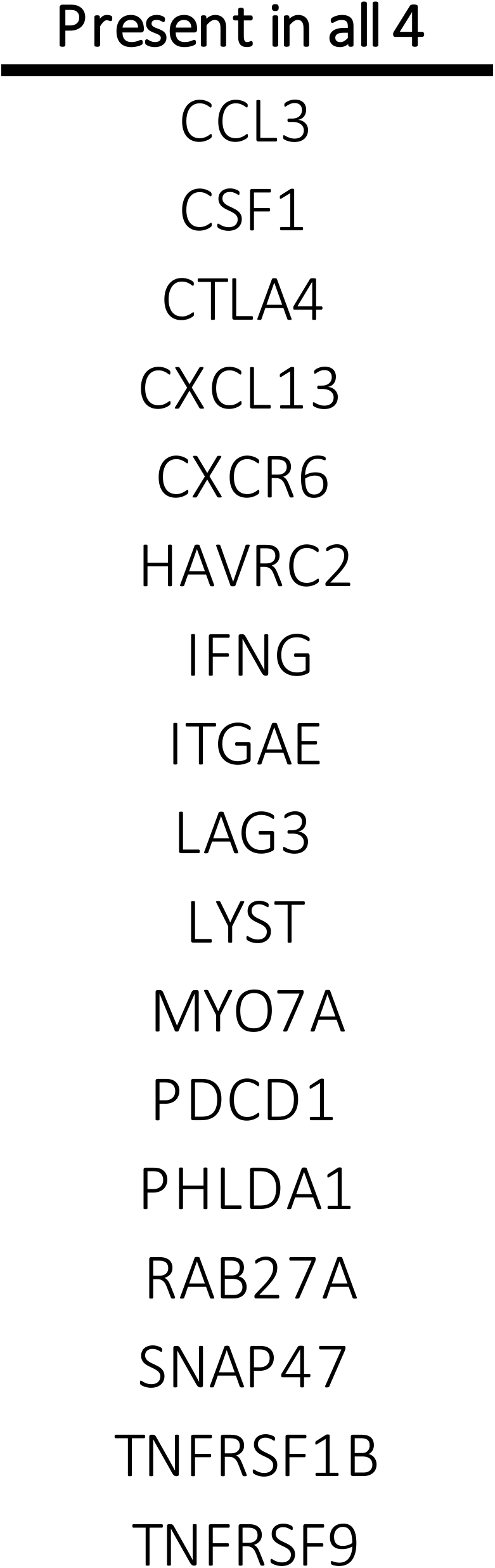
Human T cell exhaustion genesets

**Supplementary Table 3:**
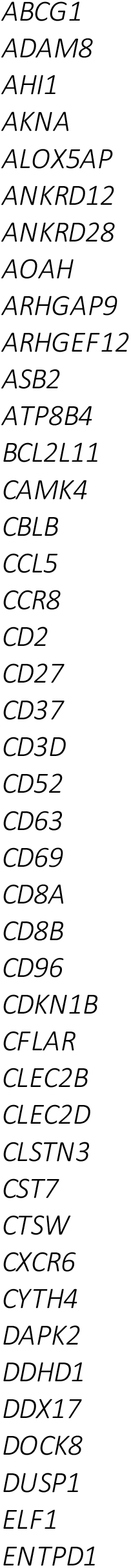
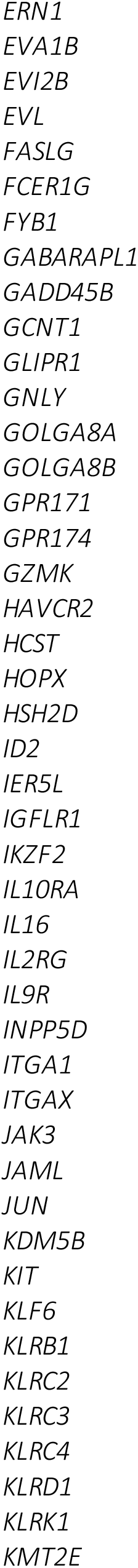
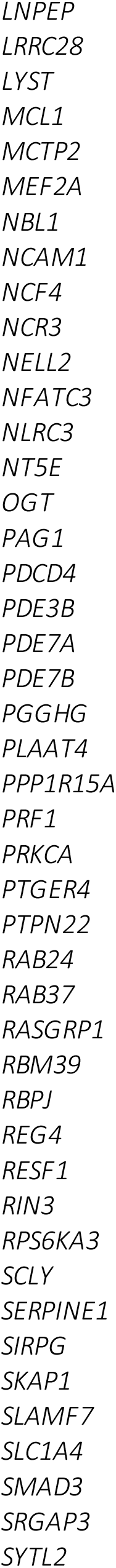
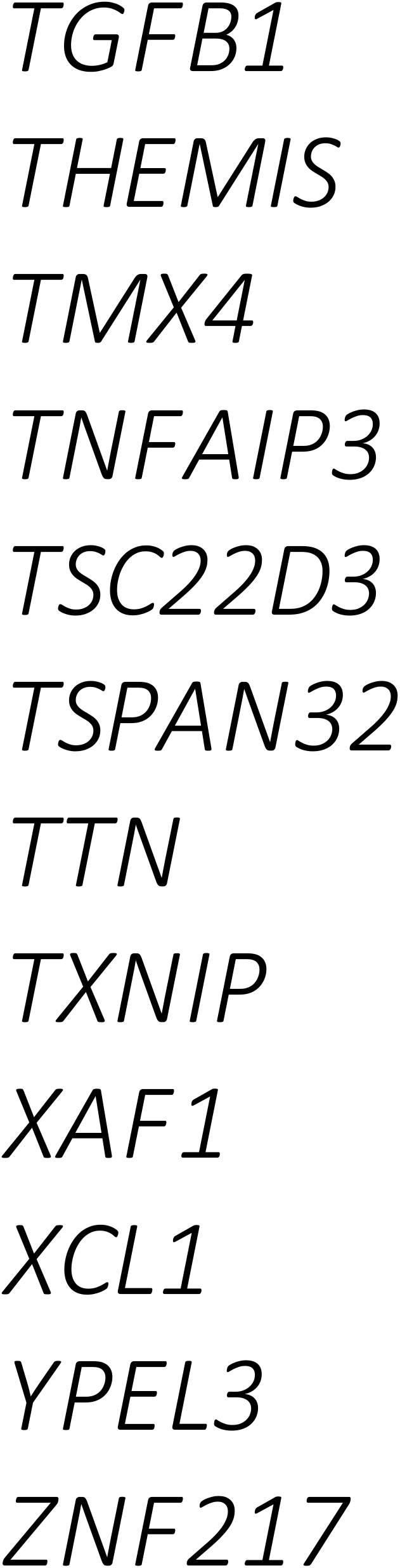
Dysfunctional 41BB-based CAR T cell (Tbbd) gene signature of dysfunction

## References

1. C. E. Brown, C. L. Mackall, CAR T cell therapy: inroads to response and resistance. Nat Rev Immunol 19, 73–74 (2019).

2. J. S. Abramson et al., Lisocabtagene maraleucel for patients with relapsed or refractory large B-cell lymphomas (TRANSCEND NHL 001): a multicentre seamless design study. Lancet 396, 839–852 (2020).

3. F. L. Locke et al., Tumor burden, inflammation, and product attributes determine outcomes of axicabtagene ciloleucel in large B-cell lymphoma. Blood Adv 4, 4898–4911 (2020).

4. L. J. Nastoupil et al., Standard-of-Care Axicabtagene Ciloleucel for Relapsed or Refractory Large B-Cell Lymphoma: Results From the US Lymphoma CAR T Consortium. J Clin Oncol 38, 3119–3128 (2020).

5. E. J. Wherry, M. Kurachi, Molecular and cellular insights into T cell exhaustion. Nat Rev Immunol 15, 486–499 (2015).

6. C. R. Good et al., An NK-like CAR T cell transition in CAR T cell dysfunction. Cell 184, 6081–6100 e6026 (2021).

7. A. H. Long et al., 4-1BB costimulation ameliorates T cell exhaustion induced by tonic signaling of chimeric antigen receptors. Nat Med 21, 581–590 (2015).

8. R. C. Lynn et al., c-Jun overexpression in CAR T cells induces exhaustion resistance. Nature 576, 293–300 (2019).

9. N. Singh et al., Impaired Death Receptor Signaling in Leukemia Causes Antigen-Independent Resistance by Inducing CAR T-cell Dysfunction. Cancer Discov 10, 552–567 (2020).

10. K. M. Cappell, J. N. Kochenderfer, A comparison of chimeric antigen receptors containing CD28 versus 4-1BB costimulatory domains. Nat Rev Clin Oncol 18, 715–727 (2021).

11. M. E. Selli, J. H. Landmann, C. Arveseth, N. Singh, Inducing T cell dysfunction by chronic stimulation of CAR-engineered T cells targeting cancer cells in suspension cultures. STAR Protoc 4, 101954 (2023).

12. O. U. Kawalekar et al., Distinct Signaling of Coreceptors Regulates Specific Metabolism Pathways and Impacts Memory Development in CAR T Cells. Immunity 44, 380–390 (2016).

13. G. J. Martinez et al., The transcription factor NFAT promotes exhaustion of activated CD8(+) T cells. Immunity 42, 265–278 (2015).

14. J. Chen et al., NR4A transcription factors limit CAR T cell function in solid tumours. Nature 567, 530–534 (2019).

15. G. Li et al., 4-1BB enhancement of CAR T function requires NF-kappaB and TRAFs. JCI Insight 3, (2018).

16. A. C. Boroughs et al., A Distinct Transcriptional Program in Human CAR T Cells Bearing the 4-1BB Signaling Domain Revealed by scRNA-Seq. Mol Ther 28, 2577–2592 (2020).

17. X. Guo et al., Global characterization of T cells in non-small-cell lung cancer by single-cell sequencing. Nat Med 24, 978–985 (2018).

18. H. Li et al., Dysfunctional CD8 T Cells Form a Proliferative, Dynamically Regulated Compartment within Human Melanoma. Cell 176, 775–789 e718 (2019).

19. L. Zhang et al., Lineage tracking reveals dynamic relationships of T cells in colorectal cancer. Nature 564, 268–272 (2018).

20. C. Zheng et al., Landscape of Infiltrating T Cells in Liver Cancer Revealed by Single-Cell Sequencing. Cell 169, 1342–1356 e1316 (2017).

21. K. E. Pauken et al., Epigenetic stability of exhausted T cells limits durability of reinvigoration by PD-1 blockade. Science 354, 1160–1165 (2016).

22. D. R. Sen et al., The epigenetic landscape of T cell exhaustion. Science 354, 1165–1169 (2016).

23. B. Bengsch et al., Epigenomic-Guided Mass Cytometry Profiling Reveals Disease-Specific Features of Exhausted CD8 T Cells. Immunity 48, 1029–1045 e1025 (2018).

24. N. D. Mathewson et al., Inhibitory CD161 receptor identified in glioma-infiltrating T cells by single-cell analysis. Cell 184, 1281–1298 e1226 (2021).

25. L. Zheng et al., Pan-cancer single-cell landscape of tumor-infiltrating T cells. Science 374, abe6474 (2021).

26. X. Qiu et al., Reversed graph embedding resolves complex single-cell trajectories. Nat Methods 14, 979–982 (2017).

27. J. J. Melenhorst et al., Decade-long leukaemia remissions with persistence of CD4(+) CAR T cells. Nature 602, 503–509 (2022).

28. S. Aibar et al., SCENIC: single-cell regulatory network inference and clustering. Nat Methods 14, 1083–1086 (2017).

29. S. M. Hedrick, R. Hess Michelini, A. L. Doedens, A. W. Goldrath, E. L. Stone, FOXO transcription factors throughout T cell biology. Nat Rev Immunol 12, 649–661 (2012).

30. Y. Harada et al., Transcription factors Foxo3a and Foxo1 couple the E3 ligase Cbl-b to the induction of Foxp3 expression in induced regulatory T cells. J Exp Med 207, 1381–1391 (2010).

31. Y. M. Kerdiles et al., Foxo transcription factors control regulatory T cell development and function. Immunity 33, 890–904 (2010).

32. A. S. Dejean et al., Transcription factor Foxo3 controls the magnitude of T cell immune responses by modulating the function of dendritic cells. Nat Immunol 10, 504–513 (2009).

33. J. Feucht et al., Calibration of CAR activation potential directs alternative T cell fates and therapeutic potency. Nat Med 25, 82–88 (2019).

34. B. Daniel et al., Divergent clonal differentiation trajectories of T cell exhaustion. Nat Immunol, (2022).

35. J. R. Giles et al., Shared and distinct biological circuits in effector, memory and exhausted CD8(+) T cells revealed by temporal single-cell transcriptomics and epigenetics. Nat Immunol, (2022).

36. Z. Jackson et al., Sequential Single-Cell Transcriptional and Protein Marker Profiling Reveals TIGIT as a Marker of CD19 CAR-T Cell Dysfunction in Patients with Non-Hodgkin Lymphoma. Cancer Discov 12, 1886–1903 (2022).

37. T. L. Wilson et al., Common Trajectories of Highly Effective CD19-Specific CAR T Cells Identified by Endogenous T-cell Receptor Lineages. Cancer Discov 12, 2098–2119 (2022).

38. J. van Grevenynghe et al., Transcription factor FOXO3a controls the persistence of memory CD4(+) T cells during HIV infection. Nat Med 14, 266–274 (2008).

39. J. A. Sullivan, E. H. Kim, E. H. Plisch, S. L. Peng, M. Suresh, FOXO3 regulates CD8 T cell memory by T cell-intrinsic mechanisms. PLoS Pathog 8, e1002533 (2012).

40. J. A. Sullivan, E. H. Kim, E. H. Plisch, M. Suresh, FOXO3 regulates the CD8 T cell response to a chronic viral infection. J Virol 86, 9025–9034 (2012).

41. N. J. Haradhvala et al., Distinct cellular dynamics associated with response to CAR-T therapy for refractory B cell lymphoma. Nat Med 28, 1848–1859 (2022).

42. Z. Good et al., Post-infusion CAR TReg cells identify patients resistant to CD19-CAR therapy. Nat Med 28, 1860–1871 (2022).

43. A. Sheih et al., Clonal kinetics and single-cell transcriptional profiling of CAR-T cells in patients undergoing CD19 CAR-T immunotherapy. Nat Commun 11, 219 (2020).

44. M. M. Berrien-Elliott et al., Hematopoietic cell transplantation donor-derived memory-like NK cells functionally persist after transfer into patients with leukemia. Sci Transl Med 14, eabm1375 (2022).

45. M. D. Robinson, D. J. McCarthy, G. K. Smyth, edgeR: a Bioconductor package for differential expression analysis of digital gene expression data. Bioinformatics 26, 139–140 (2010).

46. M. R. Corces et al., An improved ATAC-seq protocol reduces background and enables interrogation of frozen tissues. Nat Methods 14, 959–962 (2017).

47. B. Langmead, C. Trapnell, M. Pop, S. L. Salzberg, Ultrafast and memory-efficient alignment of short DNA sequences to the human genome. Genome Biol 10, R25 (2009).

48. Y. Hao et al., Integrated analysis of multimodal single-cell data. Cell 184, 3573–3587 e3529 (2021).

49. G. Finak et al., MAST: a flexible statistical framework for assessing transcriptional changes and characterizing heterogeneity in single-cell RNA sequencing data. Genome Biol 16, 278 (2015).

50. B. Van de Sande et al., A scalable SCENIC workflow for single-cell gene regulatory network analysis. Nat Protoc 15, 2247–2276 (2020).

51. T. Moerman et al., GRNBoost2 and Arboreto: efficient and scalable inference of gene regulatory networks. Bioinformatics 35, 2159–2161 (2019).

52. E. K. Brinkman, T. Chen, M. Amendola, B. van Steensel, Easy quantitative assessment of genome editing by sequence trace decomposition. Nucleic Acids Res 42, e168 (2014).

